# The meta-gut: Hippo inputs lead to community coalescence of animal and environmental microbiomes

**DOI:** 10.1101/2021.04.06.438626

**Authors:** Christopher L. Dutton, Amanda L. Subalusky, Alvaro Sanchez, Sylvie Estrela, Nanxi Lu, Stephen K. Hamilton, Laban Njoroge, Emma J. Rosi, David M. Post

**Affiliations:** Department of Ecology and Evolutionary Biology, Yale University, New Haven, CT, USA; Department of Biology, University of Florida, Gainesville, FL, USA; Microbial Sciences Institute, Yale University, New Haven, CT, USA; W.K. Kellogg Biological Station and Department of Integrative Biology, Michigan State University, Hickory Corners, MI, USA; Cary Institute of Ecosystem Studies, Millbrook, NY, USA; National Museums of Kenya, Nairobi, Kenya

**Keywords:** tropical river, hippopotamus, subsidy, anoxic, gut microbiome, community coalescence

## Abstract

All animals carry specialized microbiomes, and their gut microbiotas in particular are continuously released into the environment through excretion of waste. Here we propose the *meta-gut* as a novel conceptual framework that addresses the ability of the gut microbiome released from an animal to function outside the host and potentially alter ecosystem processes mediated by microbes. An example considered here is the hippopotamus (hippo) and the pools they inhabit. Hippo pool biogeochemistry and fecal and pool water microbial communities were examined through field sampling and an experiment. Sequencing using 16S RNA methods revealed that the active microbial communities in hippo pools that received high inputs of hippo feces are more similar to the hippo gut microbiome than other nearby aquatic environments. The overlap between the microbiomes of the hippo gut and the waters into which they excrete therefore constitutes a *meta-gut* system with potentially strong influence on the biogeochemistry of pools and downstream waters. We propose that the *meta-gut* may be present where other species congregate in high densities, particularly in aquatic environments.

**Significance:** Animals can have considerable impacts on biogeochemical cycles and ecosystem attributes through the consumption of resources and physical modifications of the environment. Likewise, microbial communities are well known to regulate biogeochemical cycles. This study links those two observations by showing that the gut microbiome in waste excreted by hippos can persist *ex-situ* in the environment and potentially alter biogeochemical cycles. This “*meta-gut*” system may be present in other ecosystems where animals congregate, and may have been more widespread in the past before many large animal populations were reduced in range and abundance.

## Introduction

Animals alter the functioning of ecosystems by consuming plant and animal matter, through the transport and excretion of nutrients, and in myriad other ways [1-6]. In many cases, the activities of animals directly or indirectly influence microorganisms, which in turn regulate the major biogeochemical cycles, although such linkages remain to be fully investigated [7-9]. Abundant evidence exists for altered decomposition and nutrient cycling in response to organic matter and nutrient excretion and egestion by animals, but few studies have explicitly examined the role of the externally released animal gut microbiome in mediating those changes [5, 6, 10-13], but see [14].

Community coalescence theory is an ecological framework for investigating the mixing of entire microbial communities and their surrounding environments [15-17], but much of the focus has been on metacommunity dynamics among the microbiota, with less attention on the ecosystem implications of resource flows that accompany this mixing [18]. Animal excretion and egestion present a unique case of community coalescence by effectively mixing the animal gut microbiome with preexisting microbial communities, together with organic matter, nutrients, and metabolic byproducts that may also be excreted or egested. Heavy rates of such loading can shape the external environment in ways that support the persistence of gut microbiota outside the host gut, particularly in aquatic ecosystems, increasing the likelihood that *ex situ* gut microbes could influence ecosystem processes and be re-ingested by other consumers. We propose that the resulting *meta-gut* system is a dynamic interplay of abiotic resources and microbial communities between the host gut, the external environment, and possibly the guts of other individual hosts that inhabit the same environment.

Hippos (*Hippopotamus amphibius*) have profound effects on aquatic ecosystems in which they wallow during the day by adding large amounts of organic matter and nutrients from their nighttime terrestrial grazing via defecation and urination [19-23], and their fecal inputs are accompanied by abundant enteric microbes (Fig. 1A). These resource subsidies interact with environmental characteristics of the recipient ecosystem to alter ecosystem function [6, 23, 24]. The organic matter excreted by hippos accumulates at the bottom of hippo pools under low to moderate discharge, and in pools with high hippo densities the decomposition of this organic matter often depletes dissolved oxygen in the water column [23]. As hippo pools become anoxic, the pool environment becomes more similar to the hippo gut, increasing the likelihood that some of the enteric microbes will survive and even function outside the host gut. Anoxia is a persistent state in the bottom waters of many hippo pools until high flow events flush organic matter downstream and reaerate the water column [23, 25], briefly exposing the microbial community to oxic conditions before organic matter loading by hippos once again drives pools towards anoxia. The conditions produced by hippos provide a unique opportunity to investigate the importance of the *meta-gut* system.

**Figure 1:**
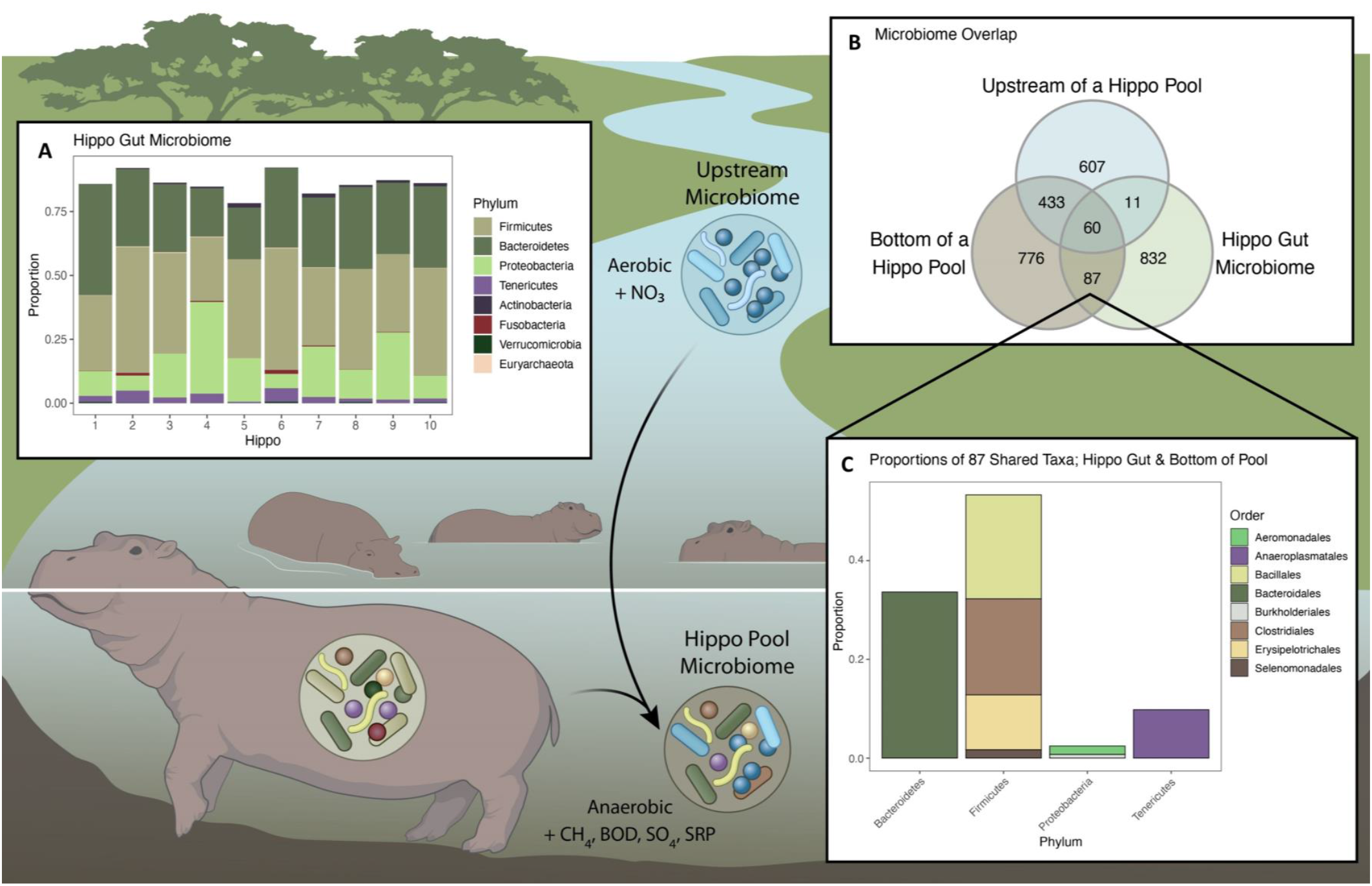
**A**. The coalescence of environmental and gut microbial communities to form the meta-gut system in a hippo pool. **B**. Proportions of the 8 most abundant phyla from the active microbial communities present in the guts of 10 individual hippos. The hippo gut microbiome was dominated by Firmicutes, Bacteroidetes and Proteobacteria. Less abundant phyla are not shown. **C**. Venn diagram showing the overlap of active microbial taxa between the hippo gut microbiome (N=10 samples), aquatic microbial communities upstream of high-subsidy hippo pools (N=19) and those from the bottom waters of high-subsidy hippo pools (N=19). **D**. Phylum and order for the 87 active taxa from the 8 most abundant orders shared between the hippo gut microbiome and the bottom waters of high-subsidy hippo pools.

The Mara River flows through the Serengeti Mara Ecosystem in Kenya and Tanzania (Fig. S1). There are over 4,000 hippos distributed among approximately 170 hippo pools in the Kenyan portion of the Mara River and its seasonal tributaries [26]. Water is present in hippo pools year-round even in the seasonal tributaries, some of which may exhibit very low or no flow during dry periods. Hippo number and water residence time of hippo pools (pool volume / discharge) interact to shape in-pool and downstream biogeochemistry in the Mara River system [23]. We classified pools by the magnitude of subsidy inputs: high-, medium-, and low-subsidy pools [38]. Low-subsidy pools remain oxic, while high-subsidy pools are typically anoxic, except during periodic flushing flows.

Here we examine this *meta-gut* phenomenon in hippo pools by sequencing the microbial communities of hippo guts and hippo pools in the field across a range of environmental conditions, and we consider the biogeochemical implications of the meta-gut system. We also conducted a microcosm experiment to investigate the role of both microbiomes and viruses (specifically bacteriophages) from the hippo gut in driving biogeochemical processes and microbial community changes within hippo pools.

## Results

### Community changes over space

We characterized the microbial communities in the hippo gut by collecting fresh hippo feces from multiple individuals across multiple pools on the landscape. There was little variability in the structure of the hippo gut microbiome among the samples from 10 individuals, and the microbiomes of all samples were dominated by Firmicutes, Bacteroidetes, Proteobacteria and Tenericutes (Fig. 1A). The gut microbiomes of hippos sharing a high-subsidy pool were not distinct relative to those of hippos from other high-subsidy pools and had similar dispersions (PERMANOVA: F = 068524, P = 0.886, PERMDISP: F = 7.3918, P = 0.013, Fig. S3).

We characterized the microbial communities in the water column of hippo pools across a gradient of hippo subsidy. Aquatic microbial communities in the bottom waters of high-, medium-, and low-subsidy hippo pools were distinctly different from one another (PERMANOVA: F = 3.3146, P = 0.002). They also had different dispersions (PERMDISP: F = 0.3532, P = 0.72); however, their differences in diversity were supported through clustering within a NMDS ordination (NMDS Stress = 0.1, Fig. 2A). The microbial communities in high-subsidy pools were more similar to those of the hippo gut microbiome than to the aquatic microbial community sampled from an area outside the influence of hippos (tributary, Fig. 2A).

**Figure 2:**
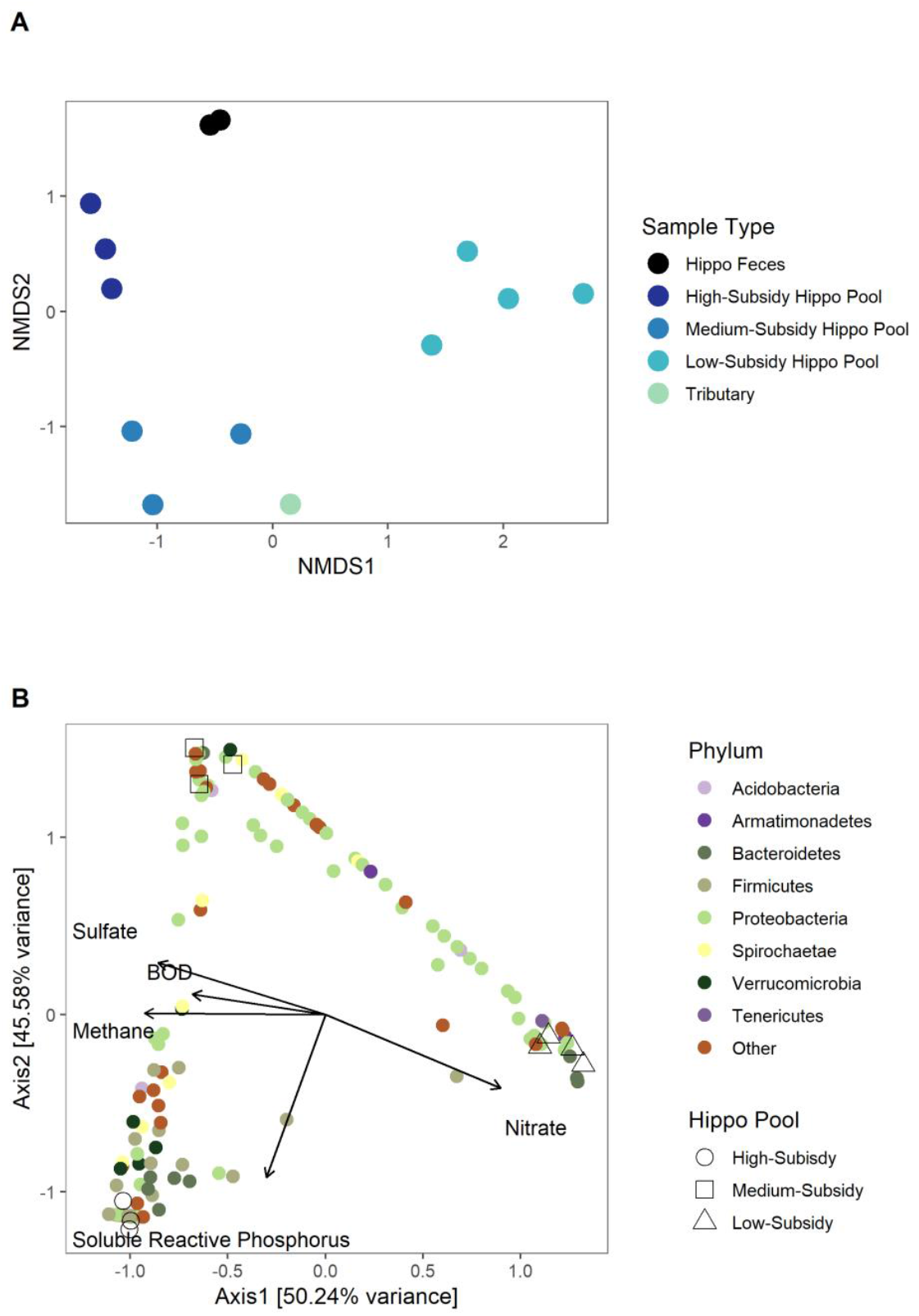
**A**. NMDS ordination (NMDS Stress = 0.1) of the Bray-Curtis dissimilarity matrix for the active aquatic microbial communities in hippo feces (N=2), in the tributary water (N=1), and in the bottom waters of high-subsidy (N=3), medium-subsidy (N=3), and low-subsidy hippo pools (N=4). **B**. Canonical correspondence analysis (CCA) of the active aquatic microbial communities (top 8 phyla are represented as colored circles with all other phyla represented as a brown circle) constrained by five environmental variables from bottom waters at the three types of sample sites: high-subsidy (hollow circle, N=3), medium-subsidy (hollow triangle, N=3), and low-subsidy hippo pools (hollow square, N=4).

We observed large differences between the three types of hippo pools when constrained by microbial communities and biogeochemistry (PERMANOVA: F = 1.3641, P = 0.011). High-, medium-, and low-subsidy hippo pools were strongly separated by the CCA ordination, which accounted for approximately 90% of the variability in the active microbial community structure within the first two axes of the constrained ordination (Fig. 2B). We observed strong relationships between the microbial community and biogeochemical constituents affected by microbial metabolism. Concentrations of dissolved methane, sulfate, and BOD were correlated and had effects (influence) that were opposite from nitrate, all of which was strongly related to axis 1 (Fig. 2B). Soluble reactive phosphorus (SRP) was strongly related to axis 2. Low-subsidy hippo pools were strongly associated with higher concentrations of nitrate. Medium-subsidy hippo pools were strongly associated with higher concentrations of sulfate, methane and BOD, but low levels of SRP. High-subsidy hippo pools were strongly associated with higher concentrations of methane, BOD, sulfate and SRP. Firmicutes were more closely associated with high-subsidy pools and higher concentrations of methane, BOD, and SRP.

We characterized the microbial communities in the water column upstream of and within hippo pools, and along a gradient of hippo density in the Mara River and its tributary, the Talek River. Aquatic microbial communities upstream of hippo pools were more variable than the communities found in the bottom waters within hippo pools (Fig. 3A). Active microbial communities in the Mara and Talek rivers were similar to one another and dissimilar to communities in hippo feces (Fig. 3B). Active microbial communities in the bottom waters of high-subsidy hippo pools were similar to both the riverine communities and the hippo feces communities. Along the upstream (site 1) to downstream (site 10) gradient of hippo influence found in the Mara River, the Mara River microbial community shifted upward on the ordination towards the microbial communities found in the Talek and in the bottom water of high-subsidy hippo pools. DO was lowest in hippo feces and the bottom of high-subsidy hippo pools. DO was highest in the middle reaches of the Talek river.

**Figure 3:**
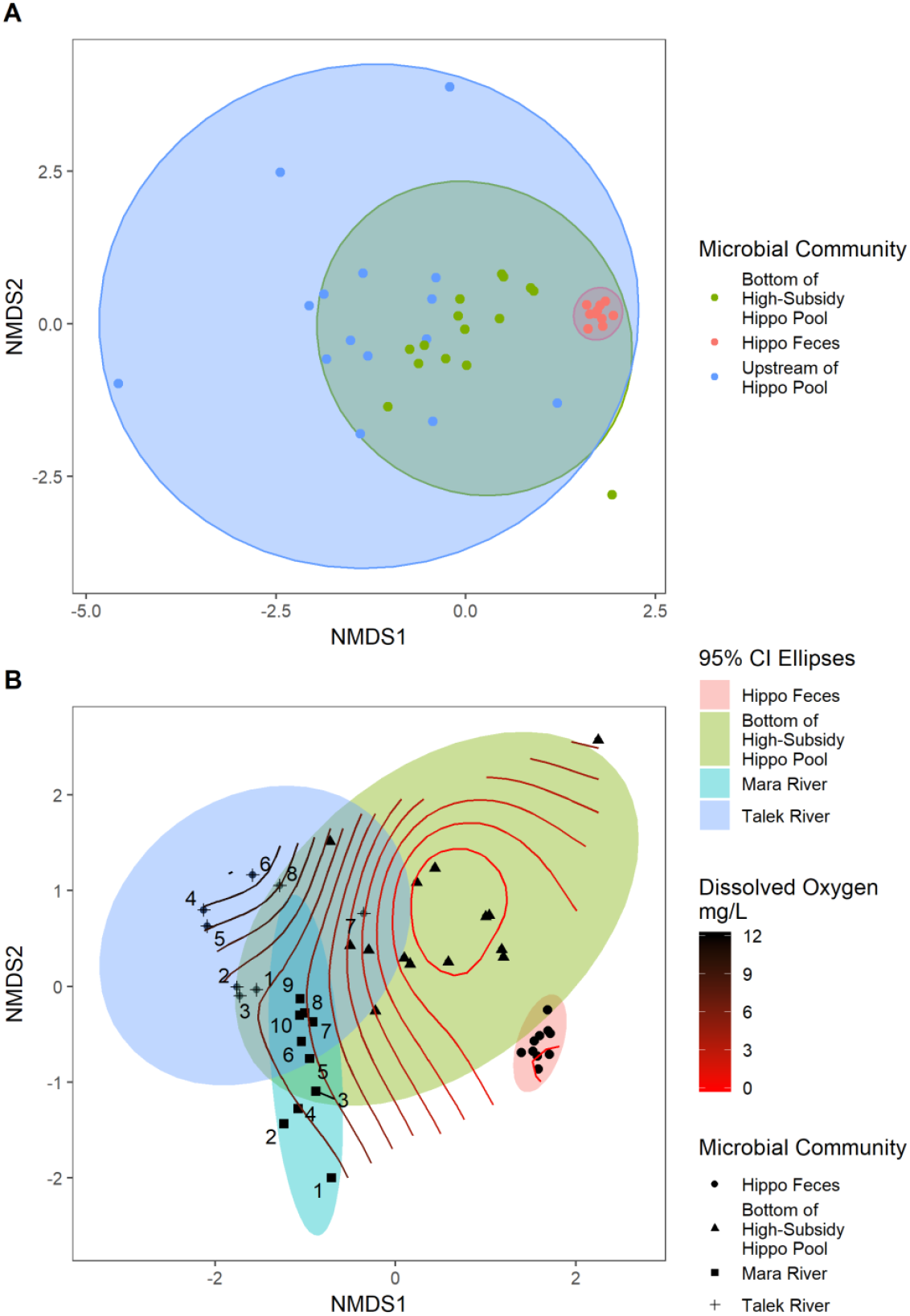
NMDS ordination of the Bray-Curtis dissimilarity matrix for the active microbial communities. **A**. Hippo feces (N=10), river waters upstream of high-subsidy hippo pools (N=15), and the bottom waters of high-subsidy hippo pools (N=15). 95% confidence ellipses are provided. NMDS Stress 0.12. **B**. Hippo feces (N=10), the bottom waters of high-subsidy hippo pools (N=15), and water samples collected along a transect of the Mara (N=10) and Talek (N=8) rivers. Numbers indicate the sampling site along the transect starting at 1 and moving downstream in ascending order. Lines represent the dissolved oxygen gradient separating the sites. 95% confidence ellipses are provided. NMDS Stress 0.12.

Eighty-seven actively functioning taxa were shared between the hippo gut microbiome (hippo feces) and the bottom waters of high-subsidy hippo pools, but not found immediately upstream of high-subsidy hippo pools (Fig. 1B). The comparatively larger overlap exhibited between the bottom of the hippo pool and the upstream sample may be due to the presence of other hippo pools located 20-100 m upstream. Most of those taxa are from the phyla Firmicutes and Bacteroidetes and are commonly found in the intestines of animals (Fig. 1C, Table S1). Notably, *Clostridia*, obligate anaerobes, were responsible for approximately 19% of the overlap between the active microbial communities in the bottom of the hippo pools and the hippo gut microbiome. *Macellibacteroides*, an obligate anaerobe, was responsible for approximately 16% of the overlap, and was previously described from anaerobic wastes from an abattoir in Tunisia [27]. Prevotellaceae, commonly found in the intestines of animals and which helps break down proteins and carbohydrates, was responsible for 11% of the overlap [28].

### Community changes over time

We characterized the microbial communities in the water column of hippo pools during transitions between aerobic and anaerobic states in response to flushing flows. As pools transitioned between anoxic and oxic states, we observed substantial changes in the proportional contribution of sources of the aquatic microbial community. In the bottom waters of all three pools, the proportion of the aquatic microbial community from the hippo gut increased after the first flushing event (7 August) until the next flushing event occurred (25 August). The increase was most pronounced in PRHP, which had over 30% of the active aquatic microbial community derived from hippo feces just prior to the August 25^th^ flushing event (Fig. 4C). Aquatic microbial communities in the bottom waters of hippo pools were more similar to those of upstream waters immediately after the flushing flow, then diverged as flows receded (Fig. 4D). In intervals between flushing flows, the aquatic microbial communities from the bottom waters of the hippo pools became more similar to hippo gut communities until the next flushing flow occurred. Across all pools, the majority of the microbial taxa could not be attributed to either hippo feces or upstream sources, which may indicate the presence of anaerobic generalists in this habitat (Figs. 4 and S4).

**Figure 4:**
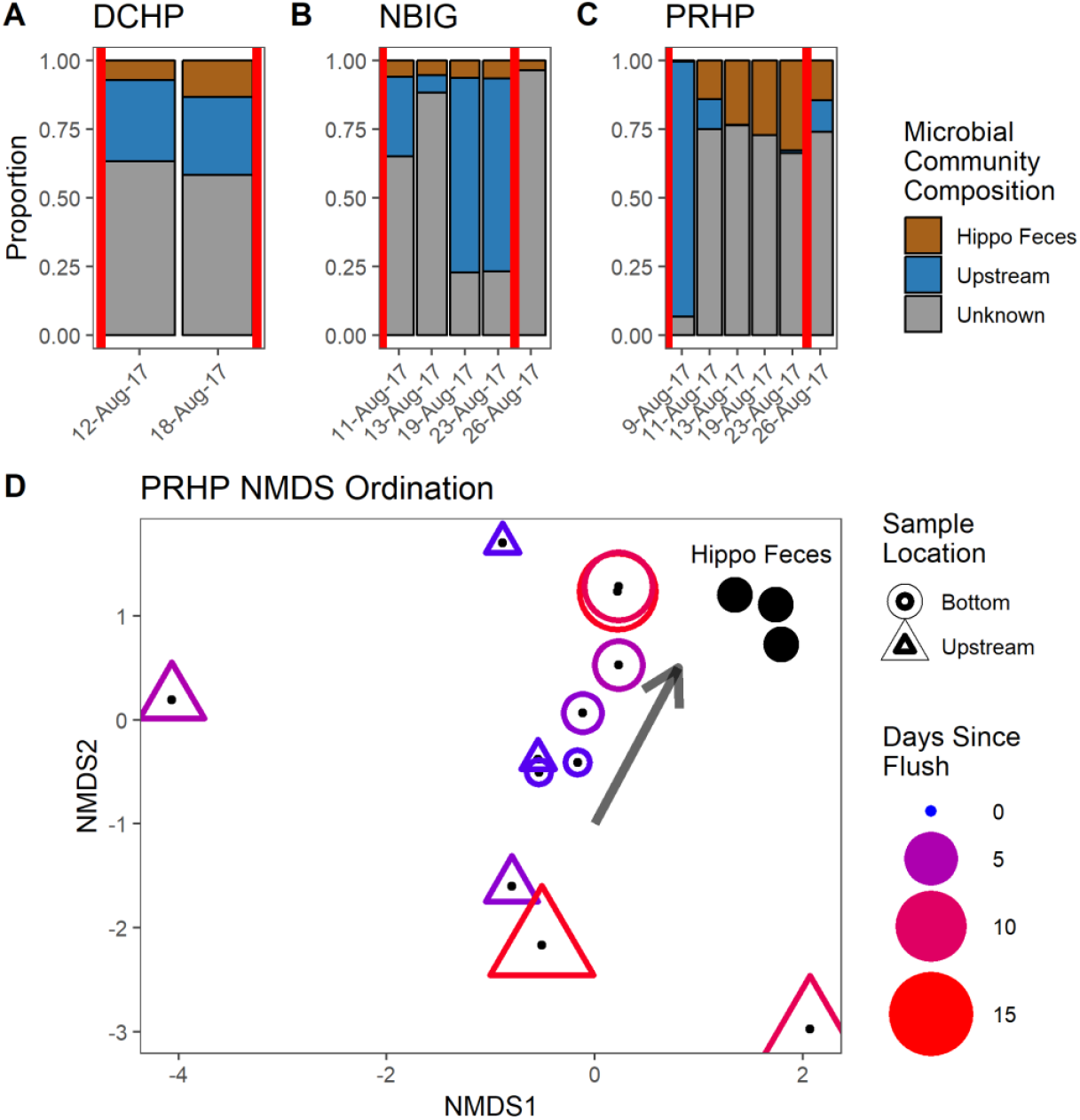
The proportion of the active microbial communities in the bottom water of high-subsidy hippo pools from hippo feces, upstream river waters, or unknown origin in **A**. DCHP Hippo Pool. **B**. NBIG Hippo Pool and, **C**. PRHP Hippo Pool. Flushing flows are represented by a red vertical line. The first flushing flow occurred on 8-Aug-17. The second flushing flow occurred on 25-Aug-17. **D**. NMDS of the Bray Curtis Dissimilatory matrix of the active microbial communities for hippo feces and the upstream and bottom samples from the PRHP high-subsidy hippo pool during the flushing events. Hippo microbiome is represented as filled black circles in the upper right of the ordination. Samples from the bottom waters are notated as a circle. Samples from the upstream are notated as a triangle. Color gradient from blue to red and size gradient from small to large represent the days since the last flushing flow. Immediately after the flushing flow, the upstream and bottom microbial communities looked similar. As time went on, the microbial community in the bottom waters looked more similar to the hippo feces microbiome than the upstream community (movement denoted as an arrow).

### Influence of hippo gut microbiome on biogeochemistry in a mesocosm experiment

We conducted a mesocosm experiment to understand the changes in microbial communities that occur as a hippo pool goes anoxic, and to test the role of microbial taxa associated with the hippo gut in driving biogeochemical changes within the hippo pools. An additional goal was to test the impact of fecal bacteriophages from the hippo gut on the microbial communities (which are composed almost entirely of bacteria) and the biogeochemical processes they mediate in hippo pools. We used a destructive sampling design with control (sterilized), bacteria, and bacteria+virus treatments (Fig. S2).

We found distinct differences in the composition of the active aquatic microbial communities through time relative to the control (Figs. 5A-D and S5). The proportion of microbes from hippo feces was largest on the first day after the start of the experiment (Day 3, September 7^th^, 2017) and declined thereafter. The total taxa (including active and inactive microorganisms) included more taxa derived from upstream than we detected in just the active taxa (Figs. 5A-C and S5).

**Figure 5:**
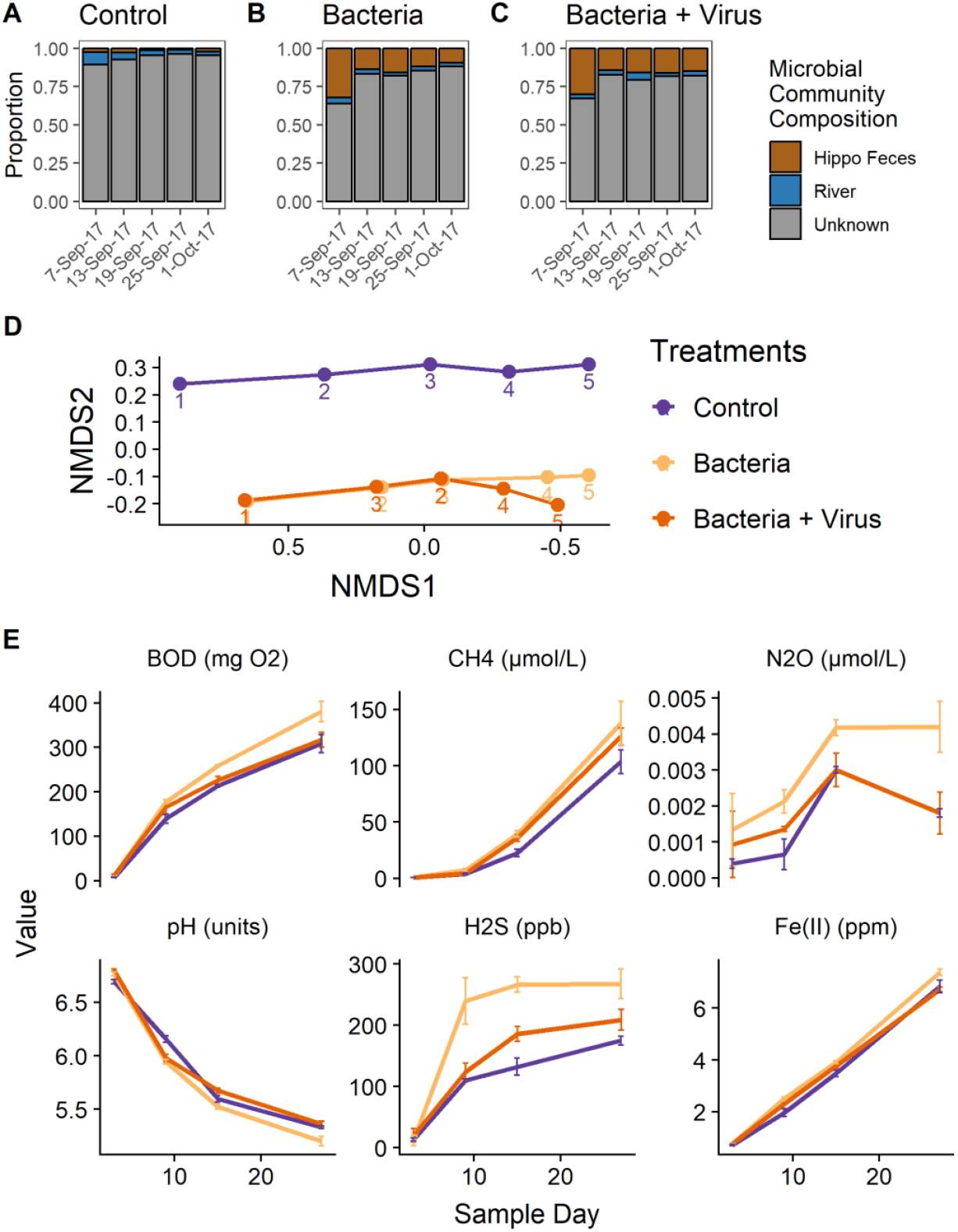
Active aquatic microbial community composition and biogeochemistry from the experimental treatments. The proportion of the active microbial community from hippo feces, upstream river water, or of unknown origin for **A**. Control. **B**. Hippo feces bacteria, no viruses. **C**. Hippo feces bacteria+virus treatments. **D**. NMDS of the Bray Curtis Dissimilatory matrix of the active aquatic microbial communities from the three experimental treatments across time. Numbers 1-5 represent the five sampling times for the experiment. NMDS Stress 0.09. **E**. Biogeochemical variables reflecting microbial catabolism, showing mean concentrations (N=3) and standard deviations as error bars (additional variables provided in Supplementary Information, Fig. S5).

We found a statistically significant effect of treatment on biogeochemical variables including Fe(II), pH, H_2_S, BOD, CH_4_, and SO_4_^2-^(Table 1). The bacteria treatment had higher concentrations of Fe(II) and lower pH compared to the bacteria+virus treatment. The bacteria treatment also had higher concentrations of BOD, H_2_S, Fe(II) and lower pH in compared to the control. There were no differences in biogeochemistry between the control and the bacteria+virus treatment.

**Table 1:** Linear mixed effect model results for each biogeochemical variable. Variables are ordered from top to bottom by statistically significant differences. Letters indicate significant differences determined by the pairwise posthoc test using Tukey’s adjusted p-value for multiple comparisons.

| Biogeochemical variable | LME - Treatment Effect |  | Pairwise Posthoc Test Treatment |  |  |
| --- | --- | --- | --- | --- | --- |
|  | F-value | p-value | Control | Bacteria | Bacteria + Virus |
| Fe(II) | 24.200 | **0.001** | **a** | **b** | **a** |
| pH | 15.999 | **0.004** | **a** | **b** | **a** |
| H<sub>2</sub>S | 11.104 | **0.010** | **a** | **b** | **ab** |
| BOD<sub>5</sub> | 9.557 | **0.014** | **a** | **b** | **ab** |
| CH<sub>4</sub> | 6.177 | **0.035** | a | a | a |
| SO<sub>4</sub><sup>2-</sup> | 5.902 | **0.038** | a | a | a |
| Chl *a*. | 4.827 | 0.056 | a | a | a |
| Cl<sup>-</sup> | 4.081 | 0.076 | a | a | a |
| SRP | 2.408 | 0.171 | a | a | a |
| NO<sub>3</sub><sup>-</sup>-N | 1.406 | 0.316 | a | a | a |
| F<sup>-</sup> | 1.339 | 0.330 | a | a | a |
| N<sub>2</sub>O | 1.232 | 0.356 | a | a | a |
| NH<sub>4</sub><sup>+</sup>-N | 1.225 | 0.358 | a | a | a |
| K<sup>+</sup> | 0.964 | 0.434 | a | a | a |
| Na<sup>+</sup> | 0.946 | 0.440 | a | a | a |
| Br<sup>-</sup> | 0.855 | 0.471 | a | a | a |
| Ca<sup>2+</sup> | 0.673 | 0.545 | a | a | a |
| DOC | 0.272 | 0.771 | a | a | a |
| Mg<sup>2+</sup> | 0.247 | 0.789 | a | a | a |
| ORP | 0.233 | 0.799 | a | a | a |
| CO<sub>2</sub> | 0.233 | 0.799 | a | a | a |

We found that the microbial communities in the bacteria and the bacteria+virus treatments were relatively similar across time until the last two sample days (sample day 23 and 27) when they began to diverge (Bray-Curtis dissimilatory matrix and NMDS ordination; Fig. 6D). Microbial communities from the last time point in the experiment consisted primarily of obligate anaerobes rather than communities associated with the hippo gut microbiome.

## Discussion

Hippos excrete microbes into the water of hippo pools, and a portion of the microbial taxa continue to function and may appreciably alter environmental biogeochemical processes (Fig. 1) [23]. The collective hippo-environmental microbiome, the *meta-gut*, is reinforced through the constant loading of hippo feces into hippo pools, which depletes dissolved oxygen and increases the similarity between the host gut and the aquatic environment.

Biogeochemistry within pools was driven by interactions between hippo loading and environmental characteristics of the pool. Microbial communities in high-subsidy hippo pools were strongly associated with higher levels of soluble reactive phosphorus (Fig. 2B), which is likely due to inputs of P in hippo urine and to the release of SRP from sediments under anoxic conditions. The oxic waters of low-subsidy hippo pools had higher concentrations of nitrate than found in the main river and there was higher sulfate found in the medium-subsidy pools in the tributaries [23]. The tributaries are likely draining catchments that have more geological sources of sulfur, while the higher nitrate concentrations in the low-subsidy hippo pools in the river may be due in part to oxic conditions that result in less denitrification. These biogeochemical differences likely interact with the rate of loading by hippos to alter microbial communities within the pools, which in turn may further alter the biogeochemistry.

We hypothesized that hippos within a pool would have more similar gut microbiomes than hippos across pools. Data from this study did not support that hypothesis—hippos from the same high-subsidy pool did not have more similar microbiomes to one another than to hippos from other high-subsidy pools (Fig. S2). However, our sample size was small (N=10, 3 feces samples per 3 pools and one additional sample near another pool), and samples were only collected near high-subsidy pools soon after a flushing flow. It is possible that similarity amongst individual gut microbiomes is more likely to develop over time between flushing flows or during dry periods when hippos congregate in larger numbers within the remaining pools.

The proportion of the active microbial community from the hippo gut in high-subsidy hippo pools increased over time between flushing flows. At most time points, the predominant source for the active microbial communities was unknown (Fig. 4A-C). Given that high-subsidy hippo pools stay anoxic for prolonged periods of time, the unknown taxa are likely anaerobic generalists that have grown in the anoxic environment in the bottom of the hippo pools and cannot be ascribed specifically to originating either from the upstream environment or the hippo gut. As hippo loading continues after a flushing flow, the hippo gut microbial community likely outcompetes certain anaerobic generalists resident in the pool while coalescing into a new community derived in part from the hippo gut microbiome. The high-subsidy pool with the highest proportion of hippo gut microbiome derived communities (PRHP) also had the highest concentrations of indicators of microbial catabolism, including BOD, NH_3_ + NH_4_^+^, SRP, DOC, CH_4_, and CO_2_ compared to the other high-subsidy pools [23]. The hippo gut microbial communities either facilitate a more reducing environment *ex situ* or thrive better within it.

Our mesocosm experiment showed that a higher proportion of active hippo gut bacteria was associated with higher concentrations of BOD, CH_4_, N_2_O, and H_2_S, suggesting that hippo gut microbes are driving some of biogeochemical differences we observed in high-subsidy hippo pools. In our experiment, the proportion of the microbial community derived from the hippo gut microbiome decreased over time, in contrast to field measurements showing an increase in gut microbes over time between flushing flows (Figs 4 & 5). This difference may be because continual inputs of hippo feces and gut microbes, which are relatively constant in high-subsidy hippo pools [29], may be necessary to outcompete other communities. It may also be that input of hippo gut microbes directly into an anoxic environment, as occurs in high-subsidy hippo pools in the field, may increase successful colonization of the pool environment. During the experiment, we loaded the hippo gut microbiome into an oxic environment at the beginning of the experiment and the bottles received no further loading. The anoxic conditions frequently present in high-subsidy hippo pools may be necessary to increase the survival of hippo gut microbes [16]. The final microbial communities in all treatments were composed of primarily generalists not derived from the hippo gut, such as *WCHB1-32* (10% average across treatments), *Acinetobacter* (9%), and Holophagaceae (6%).

Microbially-mediated transformations of biologically important elements were altered in the presence of bacteriophages from the hippo gut. The experimental treatments with a low viral load (bacteria treatment) had higher Fe(II) and lower pH than the treatments with a high viral load (bacteria+virus treatment) (Fig. 5E). We also found higher concentrations of BOD and H_2_S in the bacteria treatment than the control, with no difference between the bacteria+virus treatment and control. We hypothesize that the effect of bacteriophages could be greater when the transmission of feces occurs directly into an anoxic environment, as these conditions would decrease the presence of protists that could consume the bacteriophages [30, 31]. In both the hippo gut and in hippo pools, bacteriophages may exert particularly important control over the microbial community due to the lack of protists in anoxic environments and the likely high viral load of hippo feces [30-32]. Viral lysis of bacteria has been shown to alter biogeochemical processes such as nitrogen remineralization [33]. Our experiment results suggest that bacteriophages are inhibiting the activity of bacteria in the hippo pools, including those directly relevant to biogeochemical cycling.

Hippos subsidize tropical rivers through carbon and nutrient loading [20, 21, 23, 24, 34, 35], and here we show that they also subsidize it through loading of their gut microbiome, which uniquely ties hippos to the health of the entire riverine ecosystem. Gut microbiota provide a number of important functions that hosts are either unable to do or can do only in a limited capacity, including synthesizing amino acids, fatty acids, vitamins, and the metabolism of carbohydrates [36]. When hippos defecate into pools, many of these by-products are made directly available to aquatic life. With these by-products of gut metabolism available in the external environment (hippo pool), other aquatic life may utilize them as a subsidy [37]. Aquatic insects and fishes near hippo pools also may participate in the *meta-gut* system through the colonization of hippo gut-derived microbes into their guts. Fish gut microbiomes typically vary in response to environment and diet [38], and multiple species of fish in tropical rivers consume hippo feces [35]. If hippo gut-derived microbes are able to establish within the fish gut, they may aid in the digestion of hippo feces, or be transported to new locations by fish movements. Intriguingly, a previous meta-analysis of fish microbiomes found an unexpected similarity in fermentative bacteria between fish gut communities and vertebrate gut communities, highlighting the potential for this effect [39]. Aquatic insects also have been known either to ingest gut microbiota from the environment that can then aid directly in digestion or to rely on external microbial decomposition to process recalcitrant organic matter prior to ingestion [40].

Hippos provide an ideal case study for examining the *meta-gut* system in wild populations, as they congregate in high densities in pools that quickly become anoxic due to a subsidy overload [23, 25]. Transmission of the microbiome to the external environment may be strongest when transmission occurs directly between similar environments, such as from the gut of an animal to an anoxic aquatic environment that protects the microbiome from external stressors (atmospheric oxygen, ultraviolet radiation, adverse temperatures, etc.) [41]. This dynamic may occur broadly in other animals that wallow [42-44], share latrine sites [45], or share a common aquatic environment that can experience hypoxia/anoxia such as mussels [46, 47]. Extinct lineages of hippo-like animals or freshwater megafauna with similar environmental niches may have produced similar dynamics prior to their extinctions [48-50], suggesting that species losses of current freshwater megafauna, many of which are threatened with extinction or locally extirpated, could have even broader impacts on ecosystem functioning than previously anticipated [50-52].

## Conclusions

Altogether our research demonstrates that community coalescence between the hippo gut microbiome and the river microbiome can occur, forming a *meta-gut* system in which the hippo gut microbiota can continue to function *ex situ* in certain environmental contexts. Thus, the role of hippos in subsidizing tropical rivers extends beyond the loading of nutrients and organic matter to also include the successful transference of microbial communities that may influence biogeochemical cycling. In hippo pools, two factors are required for the *meta-gut* system. First, there is a continual loading of waste products (organic matter and nutrients) and microbial communities. Without continual loading, the resulting microbial community may coalesce into an anaerobic generalist community that does not represent the functioning gut microbial communities from the donor host. Second, loading occurs in an environmental patch that has similar environmental characteristics to the gut, which in this case develops in response to organic matter loading by hippos. Our research demonstrates that some animals can function as mobile anaerobic bioreactors, consuming organic matter, accumulating and selecting for microbes in their guts, and moving and depositing waste products and gut microbes across the landscape, where the microbes may affect ecosystem functioning of the external environment.

## Methods

### Microbial Community Sampling

#### Hippo Gut Microbiome

We characterized the microbial communities in the hippo gut by collecting ten samples of fresh hippo feces adjacent to four hippo pools early in the morning in September 2017. We collected feces from different pools and locations adjacent to the pool to include the feces of different individuals so we could estimate the similarity of gut microbiome among individuals and across the landscape. The four hippo pools are sufficiently far apart that there was likely no intermixing of hippos among them. Hippo feces from each sample was retained on a Supor polysulfone membrane, preserved in RNALater, and frozen for analysis. We assessed the composition of the microbial communities using 16S rRNA sequencing (for more details, see SI Appendix).

#### Aquatic ecosystem

We characterized the microbial communities in the water column of hippo pools across a gradient of hippo subsidy (July 2016, N=12 pools. We collected samples from the upstream, downstream, surface, and bottom of both pools containing hippos and pools that lacked hippos. Subsamples were also analyzed for biogeochemical variables (details provided below). We also collected water samples in four of the high-subsidy hippo pools every 2-3 days starting immediately after a flushing event until the next flushing event (Fig. S1) [23].

We sampled the aquatic microbial community and biogeochemical variables along a longitudinal transect down both the Mara and Talek rivers (Fig. S1, Table S1). For the Mara River, we sampled an approximately 100-km transect along a gradient of hippo numbers (N=10 locations, from 0 to ∼4,000 hippos). For the Talek River, we sampled an approximately 30-km transect to the confluence with the Mara (N=8). Mara River sites 9 and 10 are downstream of the confluence with the Talek River. Water samples were collected from each site in a well-mixed flowing section away from any hippo pools.

Aquatic microbial samples were collected by filtering water samples through a Supor polysulfone filter (0.2-µm pore size; Pall, Port Washington, NY, USA) and then preserving the filter in RNALater Stabilization Solution (Ambion, Inc., Austin, TX, USA). In 2017, the filters were preserved with RNA Later and then frozen for analysis. We used 16S rRNA sequencing of both DNA and RNA to characterize the total and active bacterial communities, respectively [53] (for more details, see Supplementary Information 1).

### Mesocosm Experiment

We collected river water from the Mara River upstream of the distribution of hippos and placed it in 45 1-L bottles in a large water basin covered by a dark tarp to help regulate temperature and prevent algal production. Bottles were randomly assigned to the control, bacteria, and bacteria+virus treatment. We collected fresh hippo feces from multiple locations adjacent to the Mara River. After homogenization, half of the hippo feces was sterilized in a pressure cooker, which testing confirmed had similar sterilization results as an autoclave [54] (see Supplementary Information 1). Five g of sterilized hippo feces was placed into each bottle to provide an organic matter substrate without viable bacteria or viruses. The unsterilized hippo feces was expressed, and the resulting liquid was filtered through 0.7-µm GF/F filters (0.7-µm pore size; Whatman, GE Healthcare Life Sciences, Pittsburgh, PA, USA) and 0.2-µm Supor filters to physically separate the bacteria (on the filter papers) from the viruses (in the filtrate). Half the filtrate was then sterilized with a UV light treatment. For more details, see Supplementary Information 1.

We prepared 15 bottles for each of three treatments—control, bacteria, and bacteria+virus—as follows: *Control* -Unfiltered river water, 5 g wet weight sterilized hippo feces, and two blank Supor filters; *Bacteria* – Unfiltered river water, 5 g wet weight sterilized hippo feces, two Supor filters containing bacteria, and 4 ml sterilized filtrate; *Virus* – Unfiltered river water, 5 g wet weight sterilized hippo feces, two Supor filters containing bacteria, and 4 ml unsterilized filtrate containing viruses.

We ran the experiment for 27 days from September to October 2017. Initial microbial samples of the river water, hippo feces bacteria and hippo fecal liquid filtrate were taken on day 0, and three replicate samples per treatment were destructively sampled on day 3, 9, 15, 21, and 27. During each time step, the bacterial communities were sampled using the methods detailed above, and chemical analyses were done on the water samples as described below. We also measured chlorophyll a, dissolved oxygen, temperature, conductivity, total dissolved solids, turbidity, and pH with a Manta2 water quality sonde (Eureka Water Probes, Austin, TX, USA).

### Chemical Analyses

All water samples collected in the field and in the experiment were analyzed for dissolved ferrous iron (Fe(II)), hydrogen sulfide (H_2_S), dissolved organic carbon (DOC), inorganic nutrients, major ions, dissolved gases, and biochemical oxygen demand following the standard methods provided in detail in Dutton et al. (2020) and briefly summarized in Supplementary Information 1.

### Statistical Analyses

We computed all statistical analyses in the R 4.0.3 statistical language in RStudio 1.3.1093 using α = 0.05 to determine significance [55, 56]. Error bars in the figures represent standard deviation of the means. All data and R code for the statistics and data treatments are provided in the Mendeley Data Online Repository (available at this preview link -https://data.mendeley.com/datasets/hrhy2fb7zn/draft?a=58671166-e96d-48f7-8d14-077a0f1bb2cc) [57].

We used the Bray-Curtis dissimilarity matrix followed by ordination with NMDS to examine differences between individual hippo gut microbiomes; between low-, medium-, and high-subsidy hippo pools; and between a gradient of hippo pools and the environment. We used a CCA to test for the influence of biogeochemical drivers. We used PERMANOVA and PERMDISP to test for significant differences between groups [58]. See Supplementary Information 1 for more details on analyses.

We compared aquatic microbial communities from the bottom of high-subsidy hippo pools, from hippo feces, and upstream of high-subsidy hippo pools (free of hippo gut microbiome influence) using the Bray-Curtis dissimilarity matrix followed by ordination with NMDS, and we quantified the proportion of taxa shared between the hippo gut microbiome and the bottom of the high-subsidy hippo pool but not present in the upstream samples.

We used SourceTracker to quantify the contribution of the hippo gut, upstream waters, or unknown sources to the active aquatic microbial communities in the bottom waters of 3 of the high-subsidy hippo pools between flushing flows [59]. We also used the Bray-Curtis dissimilarity matrix followed by ordination with NMDS to examine changes in the active aquatic microbial communities in one of the high subsidy hippo pools through time after flushing flows.

For the experiment, we calculated the Bray-Curtis dissimilatory matrix followed by ordination with NMDS for the active aquatic microbial communities over time in each of the three experimental treatments. We used SourceTracker to determine the proportion of the active aquatic microbial community in each treatment that originated from the hippo gut, the river water, or unknown sources [59]. We analyzed the biogeochemical differences among experimental treatments by fitting a linear mixed effects model for each of the biogeochemical variables throughout the experiment with the nlme package in R [60]. We fit the model with the restricted maximum likelihood method and a continuous autoregressive temporal correlation structure with sample day as the repeated factor. Treatment and time were fixed effects and individual bottles were treated as random effects. We conducted a pairwise post-hoc test with an ANOVA and the emmeans package in R to estimate marginal means with a Tukey adjusted p-value for multiple comparisons [61, 62].

## Author Contributions

Conceived of or designed study: CLD, ALS, SKH, EJR, DMP

Performed research: CLD, ALS, NL, AS, SKH, LN, EJR, DMP

Analyzed data: CLD

Wrote the paper: CLD, ALS, NL, AS, SE, SKH, LN, EJR, DMP

## Acknowledgements

Brian Heath and the Mara Conservancy hosted our experiment and provided logistical support. Geemi Paul, Ella Jourdain, James Landefield and Jordan Chancellor assisted with the sampling of hippo pools and the experiment. Bryan Yoon assisted with an experiment on DOC quality. Figure 1 conceptual diagram by Ann Sanderson. The Milwaukee County Zoo and Erin Dowgwillo provided hippo feces for an initial experiment. This research was approved by the National Council for Science and Technology in Kenya (research permit # NCST/RRI/12/1/BS-011/25) and funded by the US National Science Foundation (NSF DEB 1354053, 1354062, and 1753727). Additional funding was provided by the Yale MacMillan Fellowship for International and Area Studies (CLD), YIBS Small Grants Program (CLD), and the EEB Chair’s Fund (CLD).

## References

1. Ehrenfeld, J.G., Ecosystem Consequences of Biological Invasions, in Annual Review of Ecology, Evolution, and Systematics, Vol 41, D.J. Futuyma, H.B. Shafer, and D. Simberloff, Editors. 2010, Annual Reviews: Palo Alto. p. 59–80.

2. Mermillod-Blondin, F. and R. Rosenberg, Ecosystem engineering: the impact of bioturbation on biogeochemical processes in marine and freshwater benthic habitats. Aquatic Sciences, 2006. 68(4): p. 434–442.

3. Hooper, D.U., et al., Effects of biodiversity on ecosystem functioning: A consensus of current knowledge. Ecological Monographs, 2005. 75(1): p. 3–35.

4. Naiman, R.J., Animal Influences on Ecosystem Dynamics. BioScience, 1988. 38(11): p. 750–752.

5. Schmitz, O.J., et al., Animals and the zoogeochemistry of the carbon cycle. Science, 2018. 362(6419): p. eaar3213.

6. Subalusky, A.L. and D.M. Post, Context dependency of animal resource subsidies. Biological Reviews, 2018. 0(0).

7. Falkowski, P.G., T. Fenchel, and E.F. Delong, The microbial engines that drive Earth’s biogeochemical cycles. Science, 2008. 320(5879): p. 1034–1039.

8. Arrigo, K.R., Marine microorganisms and global nutrient cycles. Nature, 2004. 437: p. 349.

9. Flemming, H.-C. and S. Wuertz, Bacteria and archaea on Earth and their abundance in biofilms. Nature Reviews Microbiology, 2019.

10. Thrush, S.F., et al., Functional Role of Large Organisms in Intertidal Communities: Community Effects and Ecosystem Function. Ecosystems, 2006. 9(6): p. 1029–1040.

11. Vanni, M.J., Nutrient Cycling by Animals in Freshwater Ecosystems. Annual Review of Ecology and Systematics, 2002. 33: p. 341–370.

12. Atkinson, C.L., et al., Consumer-driven nutrient dynamics in freshwater ecosystems: from individuals to ecosystems. Biological Reviews, 2016.

13. Fukami, T., et al., Above-and below-ground impacts of introduced predators in seabird-dominated island ecosystems. Ecology Letters, 2006. 9(12): p. 1299–1307.

14. Drake, H.L. and M.A. Horn, As the worm turns: The earthworm gut as a transient habitat for soil microbial biomes. Annual Review of Microbiology, 2007. 61: p. 169–189.

15. Rillig, M.C., et al., Interchange of entire communities: microbial community coalescence. Trends in Ecology & Evolution, 2015. 30(8): p. 470–476.

16. Leibold, M.A., et al., The metacommunity concept: a framework for multi-scale community ecology. Ecology Letters, 2004. 7(7): p. 601–613.

17. Michel, L., M. Nicolas, and H.R. D., Meta-ecosystems: a theoretical framework for a spatial ecosystem ecology. Ecology Letters, 2003. 6(8): p. 673–679.

18. Mansour, I., et al., Application of the microbial community coalescence concept to riverine networks. Biological Reviews, 2018. 93(4): p. 1832–1845.

19. Wolanski, E. and E. Gereta, Oxygen cycle in a hippo pool, Serengeti National Park, Tanzania. African Journal of Ecology, 1999. 37(4): p. 419–423.

20. Stears, K., et al., Effects of the hippopotamus on the chemistry and ecology of a changing watershed. Proceedings of the National Academy of Sciences, 2018.

21. Subalusky, A.L., et al., The hippopotamus conveyor belt: vectors of carbon and nutrients from terrestrial grasslands to aquatic systems in sub-Saharan Africa. Freshwater Biology, 2015. 60(3).

22. Kanga, E.M., et al., Hippopotamus and livestock grazing: influences on riparian vegetation and facilitation of other herbivores in the Mara Region of Kenya. Landscape and Ecological Engineering, 2013. 9(1): p. 47–58.

23. Dutton, C.L., et al., Alternative Biogeochemical States of River Pools Mediated by Hippo Use and Flow Variability. Ecosystems, 2020.

24. Subalusky, A.L., et al., Organic matter and nutrient inputs from large wildlife influence ecosystem function in the Mara River, Africa. Ecology, 2018. 99(11): p. 2558–2574.

25. Dutton, C.L., et al., Organic matter loading by hippopotami causes subsidy overload resulting in downstream hypoxia and fish kills. Nature Communications, 2018. 9(1): p. 1951.

26. Kanga, E.M., et al., Population trend and distribution of the Vulnerable common hippopotamus Hippopotamus amphibius in the Mara Region of Kenya. Oryx, 2011. 45(1): p. 20–27.

27. Jabari, L., et al., Macellibacteroides fermentans gen. nov., sp. nov., a member of the family Porphyromonadaceae isolated from an upflow anaerobic filter treating abattoir wastewaters. International Journal of Systematic and Evolutionary Microbiology, 2012. 62(10): p. 2522–2527.

28. Wu, G.D., et al., Linking long-term dietary patterns with gut microbial enterotypes. Science (New York, N.Y.), 2011. 334(6052): p. 105–108.

29. Subalusky, A.L., et al., The hippopotamus conveyor belt: vectors of carbon and nutrients from terrestrial grasslands to aquatic systems in sub-Saharan Africa. Freshwater Biology, 2015. 60(3): p. 512–525.

30. Fenchel, T., L.D. Kristensen, and L. Rasmussen, Water column anoxia -vertical zonation of planktonic protozoa. Marine Ecology Progress Series, 1990. 62(1-2): p. 1–10.

31. Weinbauer, M.G., I. Brettar, and M.G. Höfle, Lysogeny and virus-induced mortality of bacterioplankton in surface, deep, and anoxic marine waters. Limnology and Oceanography, 2003. 48(4): p. 1457–1465.

32. Dudley, J.P., et al., Carnivory in the common hippopotamus Hippopotamus amphibius: implications for the ecology and epidemiology of anthrax in African landscapes. Mammal Review, 2016. 46(3): p. 191–203.

33. Shelford, E.J., et al., Virus-driven nitrogen cycling enhances phytoplankton growth. Aquatic microbial ecology, 2012. 66(1): p. 41–46.

34. Masese, F.O., et al., Are Large Herbivores Vectors of Terrestrial Subsidies for Riverine Food Webs? Ecosystems, 2015. 18(4): p. 686–706.

35. McCauley, D.J., et al., Carbon stable isotopes suggest that hippopotamus-vectored nutrients subsidize aquatic consumers in an East African river. Ecosphere, 2015. 6(4): p. 1–11.

36. Bull, M.J. and N.T. Plummer, Part 1: The Human Gut Microbiome in Health and Disease. Integrative medicine (Encinitas, Calif.), 2014. 13(6): p. 17–22.

37. Stephens, G.C. and R.A. Schinske, Uptake of Amino Acids by Marine INVERTEBRATES1. Limnology and Oceanography, 1961. 6(2): p. 175–181.

38. Egerton, S., et al., The gut microbiota of marine fish. Frontiers in Microbiology, 2018. 9(873).

39. Sullam, K.E., et al., Environmental and ecological factors that shape the gut bacterial communities of fish: a meta-analysis. Molecular ecology, 2012. 21(13): p. 10.1111/j.1365-294X.2012.05552.x.

40. Harris, J.M., The presence, nature, and role of gut microflora in aquatic invertebrates: A synthesis. Microbial Ecology, 1993. 25(3): p. 195–231.

41. Rocca, J.D., et al., The Microbiome Stress Project: Toward a Global Meta-Analysis of Environmental Stressors and Their Effects on Microbial Communities. Frontiers in Microbiology, 2019. 9(3272).

42. Bracke, M.B.M., Review of wallowing in pigs: Description of the behaviour and its motivational basis. Applied Animal Behaviour Science, 2011. 132(1): p. 1–13.

43. McMillan, B.R., M.R. Cottam, and D.W. Kaufman, Wallowing Behavior of American Bison (Bos bison) in Tallgrass Prairie: an Examination of Alternate Explanations. Vol. 144. 2000: SPIE. 9.

44. Gossow, H. and G. Schürholz, Social Aspects of Wallowing Behaviour in Red Deer Herds. Zeitschrift für Tierpsychologie, 1974. 34(4): p. 329–336.

45. Jordan, N.R., M.I. Cherry, and M.B. Manser, Latrine distribution and patterns of use by wild meerkats: implications for territory and mate defence. Animal Behaviour, 2007. 73(4): p. 613–622.

46. Stadmark, J. and D.J. Conley, Mussel farming as a nutrient reduction measure in the Baltic Sea: Consideration of nutrient biogeochemical cycles. Marine Pollution Bulletin, 2011. 62(7): p. 1385–1388.

47. Vaughn, C.C., S.J. Nichols, and D.E. Spooner, Community and foodweb ecology of freshwater mussels. Journal of the North American Benthological Society, 2008. 27(2): p. 409–423.

48. Boisserie, J.R., The phylogeny and taxonomy of Hippopotamidae (Mammalia: Artiodactyla): a review based on morphology and cladistic analysis. Zoological Journal of the Linnean Society, 2005. 143(1): p. 1–26.

49. Clementz, M.T., P.A. Holroyd, and P.L. Koch, Identifying Aquatic Habits of Herbivorous Mammals through Stable Isotope Analysis. PALAIOS, 2008. 23(9/10): p. 574–585.

50. Malhi, Y., et al., Megafauna and ecosystem function from the Pleistocene to the Anthropocene. Proceedings of the National Academy of Sciences, 2016. 113(4): p. 838–846.

51. He, F., et al., Freshwater megafauna diversity: Patterns, status and threats. Diversity and Distributions, 2018. 24(10): p. 1395–1404.

52. Doughty, C.E., A. Wolf, and Y. Malhi, The impact of large animal extinctions on nutrient fluxes in early river valley civilizations. Ecosphere, 2013. 4(12): p. art148.

53. Muscarella, M.E., S.E. Jones, and J.T. Lennon, Species sorting along a subsidy gradient alters bacterial community stability. Ecology, 2016. 97(8): p. 2034–2043.

54. Swenson, V.A., et al., Assessment and verification of commercially available pressure cookers for laboratory sterilization. PLOS ONE, 2018. 13(12): p. e0208769.

55. R Core Team, R: A language and environment for statistical computing. 2018, R Foundation for Statistical Computing: Vienna, Austria.

56. RStudio Team, RStudio: Integrated Development for R. 2016, RStudio, Inc.: Boston, MA.

57. Dutton, C.L., et al., Data from: The meta-gut: Hippo inputs lead to community coalescence of animal and environmental microbiomes. 2021: Mendelay Data.

58. Oksanen, J., et al., Package ‘vegan’. Community ecology package, version, 2013. 2(9).

59. Knights, D., et al., Bayesian community-wide culture-independent microbial source tracking. Nature Methods, 2011. 8: p. 761.

60. Pinheiro, J.C., et al., nlme: Linear and Nonlinear Mixed Effects Models. 2018.

61. Searle, S.R., F.M. Speed, and G.A. Milliken, Population Marginal Means in the Linear Model: An Alternative to Least Squares Means. The American Statistician, 1980. 34(4): p. 216–221.

62. Lenth, R., et al., emmeans: Estimated Marginal Means, aka Least-Squares Means. 2019.

