## Supplementary Information 1 for "The meta-gut: Hippo inputs lead to community coalescence of animal and environmental microbiomes"

**This PDF file includes:**

Figures S1 to S7

Table S1

Supplementary text

SI References

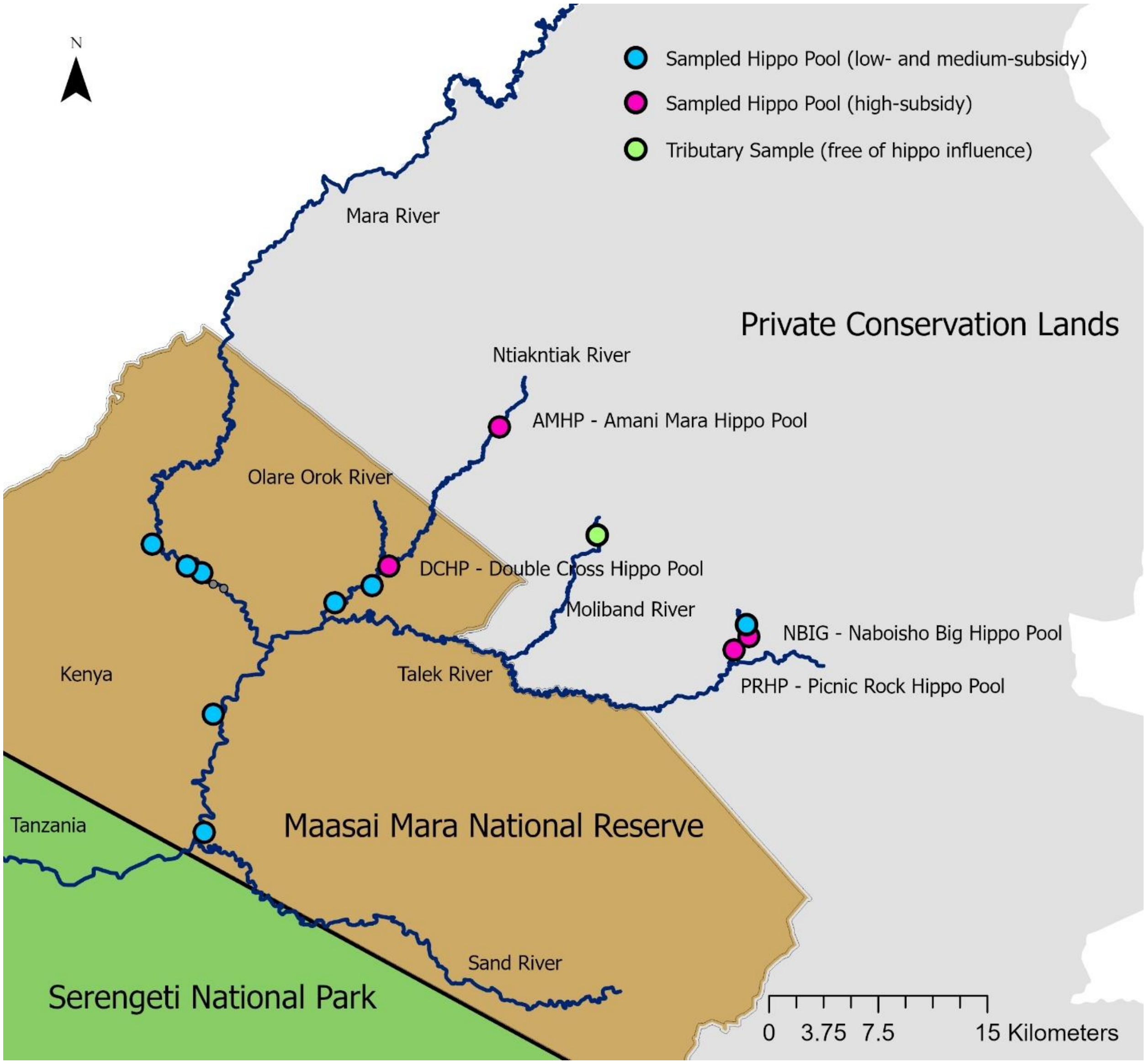


Figure S1: Hippo pools sampled across a gradient of hippo subsidy (blue and pink circles) and four high-subsidy hippo pool sampled during transitions between aerobic and anaerobic states (pink circles). One sample was collected from a tributary that was free of hippo influence (green circle).

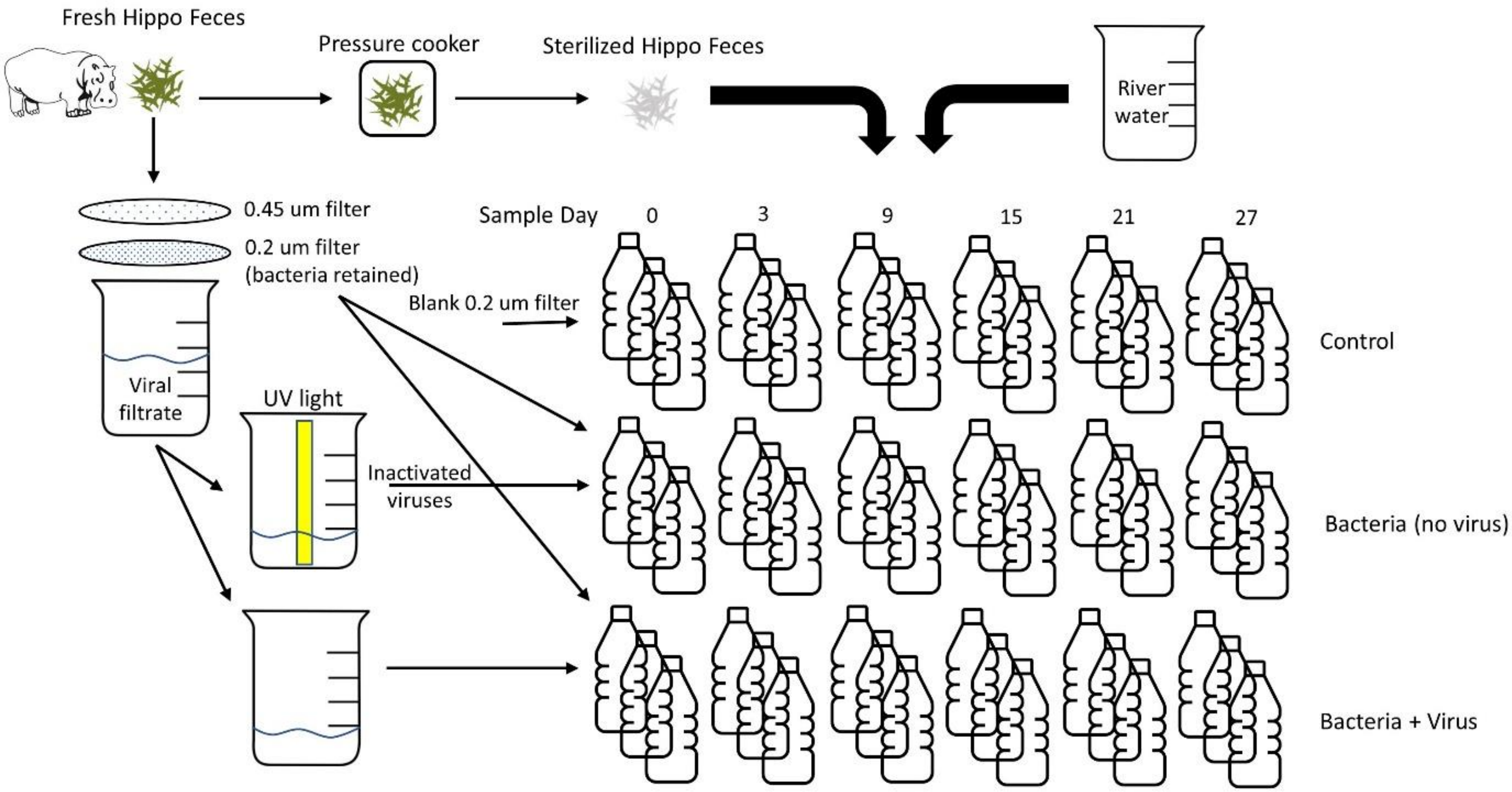


Figure S2: Microcosm experimental design to characterize the changing microbial communities as a hippo pool goes anoxic, to elucidate the role of microbial taxa associated with the hippo gut in driving biogeochemical changes, and to examine the impact of fecal bacteriophages from the hippo gut on bacterial communities and biogeochemical processes.

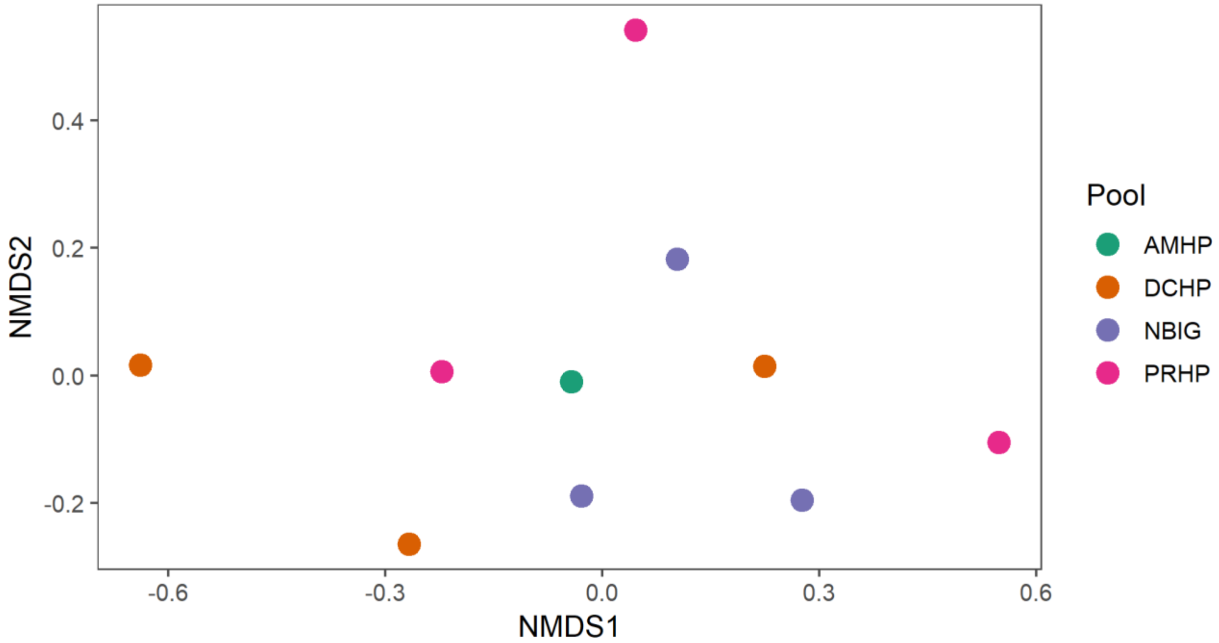


Figure S3: NMDS ordination the Bray-Curtis dissimilarity matrix of the active microbial community in hippo feces (N=10) collected near four high-density hippo pools (AMHP N=1, DCHP N=3, NBIG N=3, PRHP N=3).

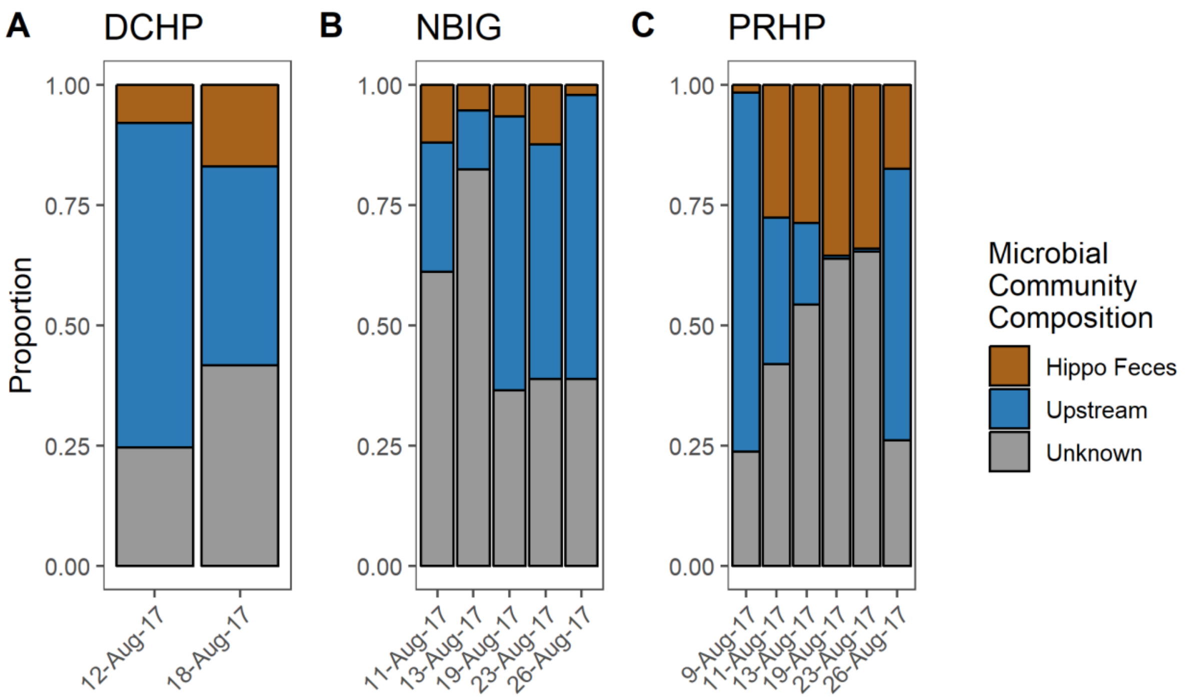


Figure S4: SourceTracker results for the total microbial communities from high-density hippo pools during the flushing flows. The first flushing flow occurred on 8-Aug-17. The second flushing flow occurred on 25-Aug-17.

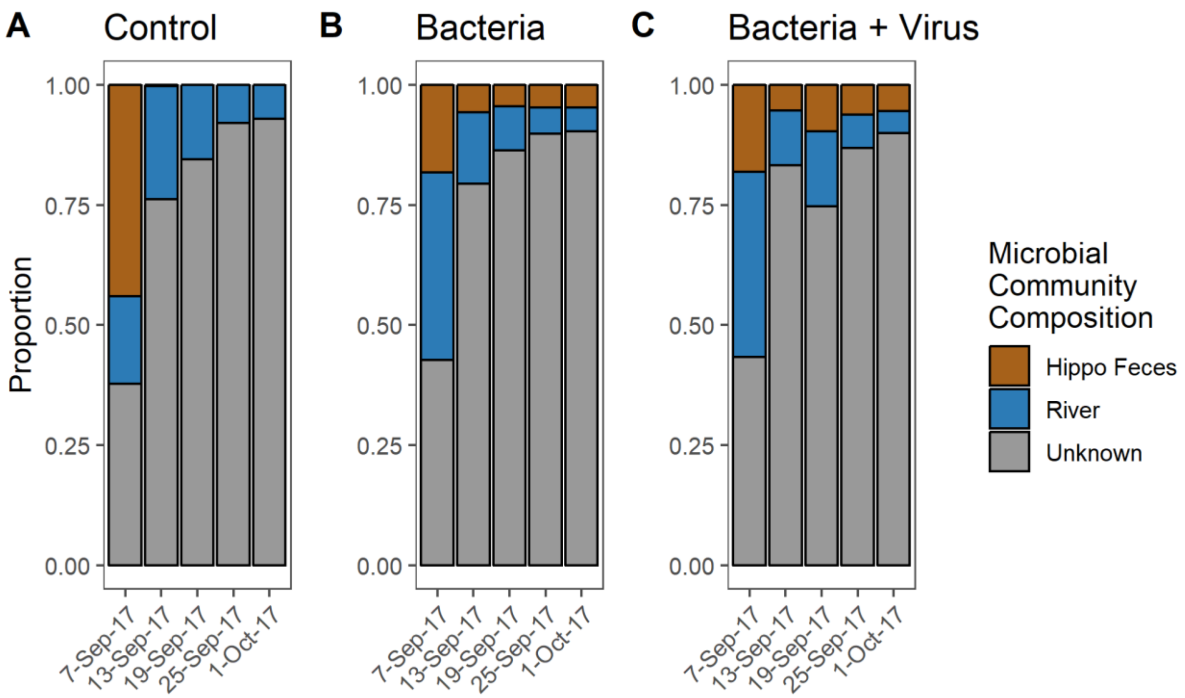


Figure S5: Total microbial community composition from the experimental treatments. SourceTracker results for **A.** Control (sterilized hippo feces only). **B**. Hippo feces plus added bacteria, no viruses. **C**. Hippo feces plus added bacteria and viruses.

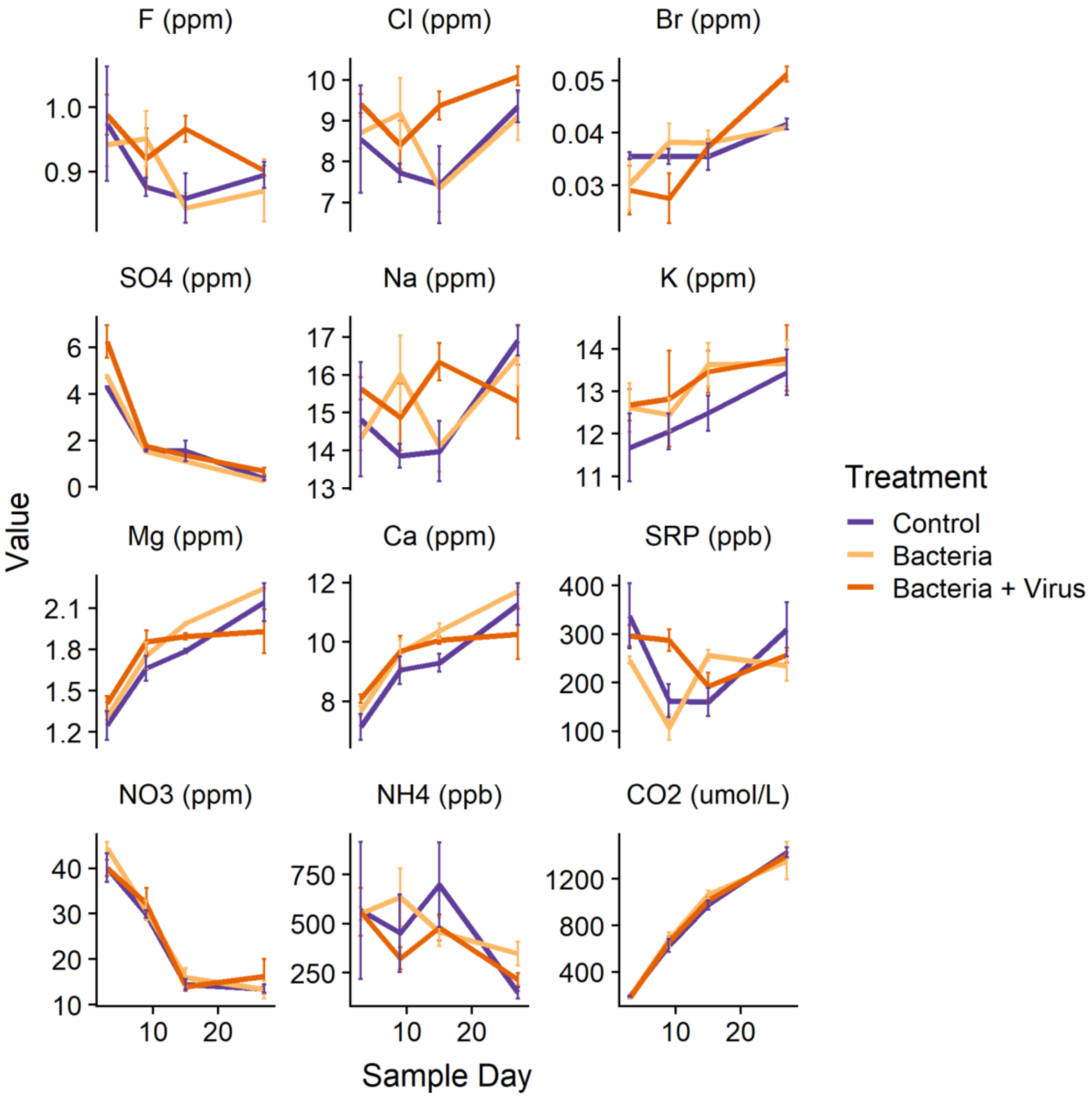


Figure S6: Additional biogeochemical variables from the experiment.

Table S1: Site locations for the longitudinal transects sampled down the Mara and Talek rivers in 2017.

| DATE | TIME | Site | Location (decimal degrees) | |
|---|---|---|---|---|
| | | | Latitude | Longitude |
| 11/8/2017 | 9:58:00 | Mara 1 | -1.05634 | 35.23141 |
| 11/8/2017 | 10:42:00 | Mara 2 | -1.09578 | 35.20228 |
| 11/8/2017 | 11:30:00 | Mara 3 | -1.13743 | 35.13311 |
| 11/8/2017 | 12:16:00 | Mara 4 | -1.22289 | 35.03641 |
| 11/8/2017 | 13:20:00 | Mara 5 | -1.31761 | 35.02211 |
| 11/8/2017 | 14:03:00 | Mara 6 | -1.39484 | 35.03452 |
| 11/8/2017 | 14:34:00 | Mara 7 | -1.43513 | 35.06147 |
| 11/8/2017 | 14:56:00 | Mara 8 | -1.46878 | 35.03342 |
| 11/8/2017 | 15:16:00 | Mara 9 | -1.50692 | 35.02615 |
| 11/8/2017 | 15:44:00 | Mara 10 | -1.54626 | 35.01909 |
| 11/4/2017 | 10:41:00 | Talek 1 | -1.47067 | 35.30169 |
| 11/4/2017 | 11:09:00 | Talek 2 | -1.46269 | 35.26452 |
| 11/4/2017 | 11:26:00 | Talek 3 | -1.45984 | 35.24873 |
| 11/4/2017 | 12:12:00 | Talek 4 | -1.44331 | 35.20824 |
| 11/4/2017 | 14:00:00 | Talek 5 | 1.42635 | 35.17826 |
| 11/4/2017 | 14:15:00 | Talek 6 | -1.41312 | 35.1353 |
| 11/4/2017 | 14:38:00 | Talek 7 | -1.41661 | 35.09698 |
| 11/4/2017 | 15:05:00 | Talek 8 | -1.42281 | 35.08352 |

### Supplementary Text

#### Microbial Sampling

##### *Hippo Pool Microbiome*

We characterized the microbial communities in the water column (also referred to as the microbiome) of hippo pools across a gradient of hippo subsidy (July 2016, N=12 pools) and during transitions between aerobic and anaerobic states in response to flushing flows (August and September 2017, N=4 pools, Fig. S1) [1]. We collected samples from the upstream, downstream, surface, and bottom of both pools containing hippos and pools that lacked hippos. Subsamples were also analyzed for biogeochemical variables (details provided below). We also collected water samples in four of the high-subsidy hippo pools (AMHP, DCHP, NBIG and PRHP) every 2-3 days starting immediately after a flushing event until the next flushing event (Fig. S1). The number of hippos, discharge and volume for each pool are presented in Dutton et al. (2020).

Aquatic microbial samples were collected by filtering water samples through a Supor polysulfone filter (0.2-µm pore size; Pall, Port Washington, NY, USA) in a polycarbonate in-line filter holder (Pall, Port Washington, NY, USA). The filter was then folded twice to preserve the biomass on the filter and stored in 14 mL of RNALater Stabilization Solution (Ambion, Inc., Austin, TX, USA).

We used a slightly altered protocol for sampling the aquatic microbial community during the flushing flow transitions (August and September 2017) in order to increase the amount of preserved genetic material in our samples. The sample was collected and filtered as before, and after the filter was dry, 15 mL of RNALater was gently poured onto the filter and the collected biomass on the filter was incubated for 15 minutes. The RNALater solution was then drawn through the filter. The filter was stored dry in a sterile petri dish and transferred to a refrigerator within several hours, then transferred to a -20 C freezer for storage within several days.

##### *Hippo Gut Microbiome*

Ten samples of the hippo gut microbiome were obtained by collecting fresh hippo feces adjacent to four hippo pools early in the morning in September 2017. We collected feces from different pools and locations adjacent to the pool to include the feces of different individuals so we could estimate the similarity of gut microbiome among individuals and from across the landscape. The four hippo pools are sufficiently far apart that there was likely no intermixing of hippos among them.

Each individual hippo feces sample was gently homogenized by hand and then the liquid was gently squeezed from the coarse particulate organic matter. A portion of the liquid (approximately 10 mL) was vacuum filtered through a Supor polysulfone membrane (0.2-µm pore size; Pall, Port Washington, NY, USA). After approximately 10 mL had filtered through and the filter was dry, 15 mL of RNALater was gently poured onto the filter and allowed to contact the collected biomass on the filter for 15 minutes before being removed by filtration. The filter was stored dry in a sterile petri dish and transferred to a refrigerator within several hours, then to a -20 C freezer for storage within several days.

During the July 2016 survey of hippo pools, we collected an additional two samples of fresh hippo feces near a high-subsidy hippo pool and filtered approximately 10 mL of the liquid portion after homogenization as detailed above. The filter was then folded twice to preserve the biomass on the filter and stored in 14 mL of RNALater.

#### Microbial Community Characterization

We used 16S rRNA sequencing to characterize the active microbial communities. We extracted both DNA and RNA from our preserved samples, then used RNA to synthesize cDNA to represent the "active" microbial community and the total DNA in the sample to represent the "total" microbes present, including those that may not be actively replicating [2]. Due to the continual loading of hippo feces into pools and the long half-life of DNA, we would expect there to be significant quantities of microbial DNA derived from hippo feces within the pools. However, there would be less accumulation of RNA because of RNA's shorter half-life. The active communities identified through this RNA-based approach are the ones that would potentially contribute to ecosystem function [3] as indicated by the protein synthesis potential, although relationships between activity and rRNA concentrations in individual taxa within mixed communities can vary [4]. Nevertheless, this method provides an overall characterization of the microbial community's potential activity.

We used the Qiagen RNeasy Powerwater Kit (Qiagen, Hilden, Germany) to extract the DNA and RNA from the material on the filter using a slightly modified manufacturer's protocol to allow for the extraction of both DNA and RNA. After extraction, we split the total extracted volume (100 µL per sample) into two groups. We treated one group with the DNase Max Kit (Qiagen, Hilden, Germany) to remove all DNA and serve as the RNA group of samples.

We used the RNA group of samples to synthesize cDNA using the SuperScript III First Strand Synthesis Kit (Invitrogen, Carlsbad, CA, USA). DNA and cDNA were quantified using the PicoGreen dsDNA Assay Kit (Molecular Probes, Eugene, OR, USA) then normalized to 5 ng/µL. Amplicon library preparation was done using a dual-index paired-end approach [5]. We amplified the V4 region of the 16S rRNA gene using dual-index primers (F515/R805) and AccuPrime Pfx SuperMix (Invitrogen, Carlsbad, CA, USA) in duplicate for each sample. We used the following thermocycling procedures:

1. 2-min initial denaturation step at 95 °C
2. 30 cycles of the following PCR
    a. 20-second denaturation at 95 °C
    b. 15-second annealing at 55 °C
    c. 5-min extension at 72 °C
3. PCR was terminated after a 10-min extension at 72 °C.

Samples were then pooled, purified and normalized using the SequelPrep Normalization Plate Kit (Invitrogen, Carlsbad, CA, USA). Barcoded amplicon libraries were then sequenced at the

Yale Center for Genome Analysis (New Haven, CT, USA) using an Illumina Miseq v2 reagent kit (Illumina, San Diego, CA, USA) to generate 2x250 base pair paired-end reads.

Sampling took place in 2016 and 2017 and involved two separate sequencing runs. The first sequencing run included negative controls and a mock community (D6306, Zymo Research, Irvin, CA, USA). The second sequencing run included negative controls, a mock community (D6306), and a single *E. coli* strain. In both runs, the mock community and single *E. coli* strain were well reconstructed from the sequences, and there was minimal contamination in the negative controls, mock community and *E. coli* strain. Analysis of the negative controls, mock community and single *E. coli* strain are provided in Supplementary Materials 2 and 3.

From those two sequencing campaigns, we received over 2 million raw sequences from the first campaign and over 7 million raw sequences for the second campaign. For the following analyses, only samples collected and sequenced during the same campaign are analyzed together to prevent preservation or sequencing biases. However, samples within the two separate campaigns were preserved and sequenced using identical methods with only a minor modification (mentioned above) to increase the preservation of genetic material.

We de-multiplexed sequenced reads then removed barcodes, indexes, and primers using QIIME2 [6]. We used DADA2 with a standard workflow in R [7] to infer exact sequence variants (ESV) for each sample [8]. We assigned taxonomy using a naïve Bayesian classifier and the SILVA training set v. 128 database [9, 10]. We removed potential contamination in samples from both campaigns by using the statistical technique in the R package, d*econtam* [11]. We used Phyloseq to characterize, ordinate, and compare microbial communities [12] with their standard workflow [7].

#### Mesocosm Experiment

We conducted a microcosm experiment to understand the changes in microbial communities that occur as a hippo pool goes anoxic, and to test the role of microbial taxa associated with the hippo gut in driving biogeochemical changes within the hippo pools. An additional goal was to test the impact of fecal bacteriophages from the hippo gut on the microbial communities (which are composed almost entirely of bacteria) and biogeochemical processes they mediate in hippo pools. We used a destructive sampling design with control (sterilized), bacteria, and bacteria+virus treatments (Fig. S2).

River water was collected from the Mara River upstream of the distribution of hippos and transported to the field camp in a plastic water container filled completely to eliminate air bubbles. The water was then placed in 45 1-L bottles laid on their side in a large water basin covered by a dark tarp to help regulate temperature and prevent algal production. Bottles were randomly assigned to the control, bacteria, and bacteria+virus treatment.

Fresh hippo feces were collected from several different locations adjacent to the Mara River. After homogenization, half of the hippo feces was sterilized in a pressure cooker (01264 6-Quart Aluminum Pressure Cooker, National Presto Industries, Eau Claire, WI, USA). Previously we compared the effectiveness of hippo feces sterilization using a pressure cooker or an autoclave, and found that pressure cooking hippo feces for 5 minutes had a similar effect as using an autoclave for one hour (see Supplementary Material 1). Similar results using pressure cookers have recently been confirmed by others [13].

For the sterilized control treatment, we added 5 g wet weight of sterilized hippo feces to each bottle to provide organic matter substrate without viable bacteria or viruses. The control treatment contained unfiltered river water, the sterilized hippo feces, and two blank Supor filters. The remainder of the non-sterilized hippo feces was used for the bacterial and bacteria+virus treatments, with each bottle receiving 5 g wet weight of feces.

The bacteria and bacteria+virus treatments were produced from the liquid expressed from the feces. The resultant cloudy liquid was diluted 10 fold with DI water and filtered through a GF/F glass fiber filter (0.7-µm pore size; Whatman, GE Healthcare Life Sciences, Pittsburgh, PA, USA) to produce a filtrate containing a bacteria+virus inoculum. We further filtered 4 mL of that liquid through multiple 0.2-µm Supor filters into a sterile glass bottle to remove the majority of but allow suspended viruses (i.e., those not associated with larger particulate matter) to pass through with the filtrate. Half of the 0.2-µm filtrate containing viruses was retained and half was treated with a 2-minute UV light treatment using a Steripen (Steripen Classic 3 UV Water Purifier, Katadyn Group, Minneapolis, MN, USA) to inactivate the viruses. We assessed whether the short-term UV light treatment would significantly alter the quality of dissolved organic matter between the treatments and determined that any change in quality was minimal and would not affect the treatments (see Supplementary Materials). To create the bacteria treatment, we placed two of the Supor filters (which would have contained particulate matter in the 0.45-0.7-µm size range) into 4 ml of inactivated filtrate and added that to the sterilized hippo feces. To create the bacteria+virus treatment, we added two Supor filters and 4 ml of untreated filtrate to the sterilized hippo feces. The bacteria treatment is not virus-free, as viruses could be attached to or infecting bacteria, but it is expected to be relatively enriched in bacteria and depleted of viruses.

We sampled our three treatments five times over 27 days in September to October 2017. All bottles were prepared at day 0. Samples of the starting constituents (river water, hippo feces bacteria and hippo fecal liquid filtrate) were taken on day 0. Three replicate samples per treatment were destructively sampled on day 3, 9, 15, 21, and 27. During each time step, the bacterial communities were sampled using the methods detailed above, and concentrations of inorganic and total nitrogen and phosphorus, dissolved organic carbon (DOC), dissolved gases ($CH_4$, $N_2O$, $CO_2$), dissolved total $H_2S$, dissolved Fe(II), $F^-$, $Cl^-$, $Br^-$, $SO_4^{2-}$, $Na^+$, $K^+$, $Mg^+$, and $Ca^{2+}$ were measured using the procedures outlined in the SI. We also measured chlorophyll a, dissolved oxygen, temperature, conductivity, total dissolved solids, turbidity, and pH with a Manta2 water quality sonde (Eureka Water Probes, Austin, TX, USA).

#### Chemical Analyses

Upon water collection for both field and experimental samples, we filtered subsamples for the analysis of dissolved ferrous iron (Fe(II)) through a 0.2 µm Supor polysulfone membrane filter directly into a solution of 50 mM 4-(2-hydroxyethyl)-1-piperazineethanesulfonic acid buffer containing ferrozine (1 g $L^{-1}$). We measured absorbances on a field spectrophotometer (DR 1900 Portable Spectrophotometer, Hach Company, Loveland, Colorado) on the day of collection. The Fe(II) analysis is based on a method modified from Lovley and Phillips [14] and Stookey [15].

Also upon collection, we filtered subsamples for the analysis of dissolved hydrogen sulfide ($H_2S$) through a Whatman GF/F glass fiber filter into two scintillation vials, taking care to avoid gas exchange, and immediately added reagents in the field following the methylene blue method of Golterman [16]. Absorbances were measured on a field spectrophotometer on the day of collection.

We filtered subsamples for dissolved organic carbon (DOC) through a pre-combusted, pre-weighed Whatman GF/F glass fiber filter and acidified the filtrate with sulfuric acid to pH <2. Analysis took place in the laboratory on a Shimadzu high-temperature, platinum-catalyzed total organic carbon analyzer (Shimadzu, Kyoto, Japan).

We preserved water samples for analysis of inorganic nutrients by filtering through a Supor 0.2-µm polysulfone membrane filter directly into a collection bottle and then freezing the filtrate. Samples were thawed to ambient temperature and analyzed on a portable flow injection analyzer in the field. Ammonium ($NH_3$ + $NH_4^+$) was analyzed using the gas exchange method [17]. Nitrate ($NO_3^-$) was analyzed using zinc reduction [18]. Soluble reactive phosphorus (SRP) was analyzed using the molybdate blue method [19].

We preserved samples for total nitrogen (TN) and total phosphorus (TP) by decanting unfiltered sample water into the collection bottle and acidifying it with sulfuric acid to pH <2. Analyses were run on an Astoria-Pacific flow analyzer in the laboratory using an alkaline potassium persulfate digestion (Astoria-Pacific, Clackamas, Oregon, USA).

Samples for analysis of major ions were preserved by filtering through a Whatman GF/F 0.45µm filter directly into the collection bottle. Analyses were run in the laboratory on a Dionex ion chromatograph system equipped with membrane suppression and conductivity detection (Dionex, Sunnyvale, California, USA)

We extracted dissolved gases from water samples by static headspace equilibration [20]. In short, 115 mL of water was drawn into a 140 mL syringe with a stopcock, and 25 mL of ambient air from upwind of hippo pools was then drawn into the syringe. The syringe was then gently shaken for 5 minutes to equilibrate the dissolved gases with the headspace, then 15 mL of headspace air was injected into evacuated exetainers and stored within water filled 50 mL Falcon tubes or a water filled 1-liter Nalgene bottle. We analyzed preserved samples of dissolved gases in the laboratory on a Shimadzu GC-2014 gas chromatograph (Shimadzu, Kyoto, Japan) in the USA. Extended travel time and several flights were needed to get the samples back to the laboratory from the field. Trip standards were utilized to account for any leakage in the exetainers due to pressure changes.

We determined the biochemical oxygen demand over five days (BOD5) for the water samples by incubating a subsample diluted to 300 mL with a phosphate buffer, and solutions containing magnesium sulfate, calcium chloride and ferric chloride [17]. Samples were incubated for approximately 24 hours in 300 mL air-tight glass bottles. We measured dissolved oxygen at the beginning and end of the incubation with a YSI ProODO optical dissolved oxygen sensor (YSI Incorporated, Yellow Springs, Ohio, USA). The change in dissolved oxygen (mg $L^{-1}$) was then used to calculate the biochemical oxygen demand over 5 days.[1]

#### Statistical Analyses

We computed all statistical analyses in the R 4.0.3 statistical language in RStudio 1.3.1093 using $\alpha = 0.05$ to determine significance [21, 22]. All R code for the statistics and data treatments are provided in Supplementary Materials 2 and 3. Error bars in the figures represent standard deviation of the means.

We characterized the gut microbiome of different individuals inhabiting different hippo pools by calculating the Bray-Curtis dissimilarity matrix on the relative abundances for the active microbial communities from hippo feces (N=10) collected next to four high-subsidy hippo pools. We used a permutational multivariate analysis of variance (PERMANOVA) with 999 permutations to test for significant differences in the individual gut microbiomes of hippos between pools [23]. The PERMANOVA test is sensitive to different dispersions within groups so we then used a permutational analysis of multivariate dispersions (PERMDISP) to test if the differences in diversity in individual gut microbiomes between pools were potentially due to differences in dispersion among hippo pools [23].

We characterized differences in aquatic microbial communities by pool type using the Bray-Curtis dissimilarity matrix on the relative abundances for the active microbial communities from high- (N=3), medium- (N=3), and low-subsidy hippo pools (N=4), from hippo feces (N=2), and from the main tributary (N=1, Talek). We then ordinated the dissimilarity matrix using a NMDS. We recalculated the Bray-Curtis dissimilarity matrix using only the three types of hippo pools (high-, medium-, and low-subsidy hippo pools) and used a PERMANOVA and PERMDISP to compare the three types of hippo pools. We then used a canonical correspondence analysis (CCA) on the relative abundances to model the aquatic microbial communities constrained by several log-transformed environmental variables from the pools. We constrained the CCA ordination by soluble reactive phosphorus, nitrate, methane, BOD, and sulfate, which were all previously shown to be important drivers in the variation between pools [1]. We then ran a PERMANOVA to test for the significance of the constrained axes used in the ordination.

We compared aquatic microbial communities from the bottom of high-subsidy hippo pools (N=15), from hippo feces (N=10, the hippo gut microbiome) and upstream of high-subsidy hippo pools (N=15, free of hippo gut microbiome influence) using the Bray-Curtis dissimilarity matrix on the relative abundances for the active aquatic microbial communities collected from the different sample types followed by ordination with NMDS. 95% confidence ellipses were generated. We then determined the active taxa that were shared between the hippo gut microbiome (hippo feces) and the bottom of the high-subsidy hippo pools and not present in the upstream samples from high-subsidy hippo pools.

We compared aquatic microbial communities from the bottom of high-subsidy hippo pools (N=15), from hippo feces (N=10, the hippo gut microbiome) and from a longitudinal transect of the Mara (N=10) and Talek (N=8) rivers using the Bray-Curtis dissimilarity matrix on the relative abundances for the active microbial communities ordinated with NMDS. 95% confidence ellipses were generated. A gradient of dissolved oxygen was generated and overlain on the ordination.

We used SourceTracker to quantify the contribution of different potential sources to the active aquatic microbial communities in 3 of the high-subsidy hippo pools between flushing flows [24]. SourceTracker was originally developed to determine the proportion of contaminants from different source environments. We used it to determine the proportion of the active aquatic microbial community in the bottom waters of hippo pools that originated from the hippo gut or upstream, the two primary sources of microbial communities. For each time point in each hippo pool, we used the active community from the upstream sample and the active community from the hippo feces collected near the pool as potential sources. The active community from the bottom of the hippo pool was input into the model as the “sink”. SourceTracker then provides estimates of the proportion of each source to the composition of the sink, including an estimate of the proportion of microbes from any unsampled (unknown) sources. To view changes in the active aquatic microbial communities after flushing flows through time, we calculated the Bray-Curtis dissimilarity matrix on the relative abundances for upstream and bottom aquatic microbial communities in one of the high-subsidy hippo pools (PRHP) and then ordinated with NMDS.

For our experimental data, we used SourceTracker to determine the proportion of the active aquatic microbial community in each treatment that originated from the hippo gut or the river, the two primary sources of microbial communities [24]. For each timepoint, we used the active hippo gut community from feces and the active aquatic community from the river collected at Emarti as potential sources. The active community from the bottle was input into the model as the “sink.” We also calculated the Bray-Curtis dissimilatory matrix on the relative abundances of the active aquatic microbial communities over time in the three experimental treatments then ordinated with NMDS.

We analyzed the biogeochemical differences among experimental treatments by fitting a linear mixed effects model for each of the biogeochemical variables throughout the experiment with the nlme package in R [25]. We fit the model with the restricted maximum likelihood method and a continuous autoregressive temporal correlation structure with sample day as the repeated factor. Treatment and time were fixed effects and individual bottles were treated as random effects. We conducted a pairwise post-hoc test with an anova and the emmeans package in R to estimate marginal means with a Tukey adjusted p-value for multiple comparisons [26, 27].

#### Preliminary experiment to test the efficacy of sterilizing hippo feces in a pressure cooker

We used sterilized hippo feces as an organic matter substrate for our bottle experiment. Because our experiment took place at our remote field camp in the Maasai Mara National Reserve, alternatives to an autoclave were explored as a method to sterilize fresh hippo feces. We tested three methods of sterilization that we would have access to at our field site; a pressure cooker, drying in the sun and drying in an oven. We used 5 replicates of each of the three treatments in addition to also using an autoclave. After attempting sterilization using the different methods, we tested biological activity by measuring dissolved oxygen over time after submerging the sterilized material in water.

**Setup**

Autoclave - We placed 15 mg of fresh hippo feces into 5 acid washed beakers. The 5 beakers of fresh hippo feces were then placed inside of an autoclave bag and loosely sealed. The autoclave was run one hour on the standard cycle.

Pressure Cooker – We placed 15 mg of fresh hippos feces into a pressure cooker (6-quart aluminium, National Presto Industries, Inc., Eau Claire, WI, USA) with a small amount of water. We applied heat to each of the five replicates in the pressure cooker and attained maximum pressure for 5 minutes.

Oven Dried – We placed five replicates of 15 mg of fresh hippo feces in a drying oven at 60C for 24 hours.

Sun Dried – We placed five replicates of 15 mg of fresh hippo feces on a sheet of aluminium foil in the sun for 6 hours on a cloudy day.

We then took each replicate from each treatment and spiked it into 200 mL of oxygenated nanopure water (8.25 mg/L). We measured dissolved oxygen in each treatment replicate after the addition at 15 minutes, 1020 minutes and 2460 minutes. We also spiked five replicates of 15mg of fresh hippo feces into 200 mL of oxygenated water.

**Results/Conclusions**

Autoclaved hippo feces and pressure-cooked hippo feces responded similarly (Fig. S7). There was a slow drop in dissolved oxygen over the experiment that likely indicates a similar amount of biological activity between autoclaving and pressure cooking. Oven drying and sun drying also had similar results, but were less effective than autoclaving or pressure cooking. Dissolved oxygen reached zero mg/L by 2460 minutes in every treatment (except for the water-only control). The water control showed no biological activity (no drop in dissolved oxygen). Untreated fresh hippo feces showed the quickest drop in dissolved oxygen.

This data does not accurately represent the slope of the oxygen decline in the hippo feces, sun dried or oven dried treatments. The dissolved oxygen was likely depleted in the hippo feces treatment within approximately 60 minutes. The dissolved oxygen was likely depleted in the sun dried and oven dried treatments within 120 minutes.

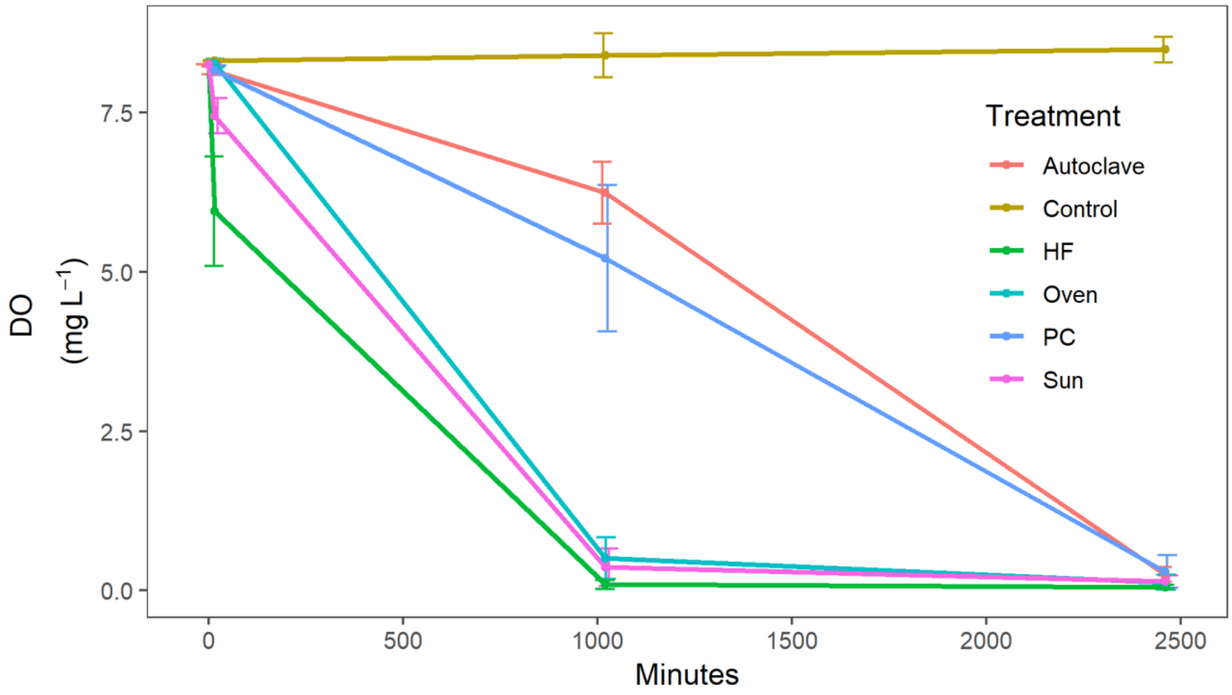


Figure S7: Dissolved oxygen measured over time for the three treatments (oven dried, sun dried, pressure cooker) as well as fresh hippo feces, autoclaved hippo feces and a control (water blank).

#### Experiment to test if a short-term UV treatment affects DOM quality

UV light can alter DOC quality and affect biogeochemical cycling. To determine if DOC quality is altered after a short 2-minute UV light treatment with a Steripen, we conducted a small experiment.

The original hippo feces liquid used in the experiment had a DOC concentration of 247 mg/L. We recreated a similar solution in the laboratory at Yale University with hippo feces from the Milwaukee County Zoo. We subjected half of that solution to a 2-minute UV light treatment and left the other half as a control. We then did dilutions using MQ water down to approximately 2 mg/L of DOC for the treatment and control. DOC quality analysis was done on an Aqualog (Horiba Scientific, Kyoto, Japan). We had two replicates of the control and of the treatment.

In the UV light treatment, we degraded aromaticity (SUVA) between 0.7 to 1.5% of the total DOM pool within each bottle and do not believe that this would cause any difference between treatments.

### Supplementary Materials References


1. Dutton, C.L., et al., *Alternative biogeochemical states of river pools mediated by hippo use and flow variability.* Ecosystems, 2020.
2. Muscarella, M.E., S.E. Jones, and J.T. Lennon, *Species sorting along a subsidy gradient alters bacterial community stability.* Ecology, 2016. **97**(8): p. 2034-2043.
3. Jones, S.E. and J.T. Lennon, *Dormancy contributes to the maintenance of microbial diversity.* Proceedings of the National Academy of Sciences, 2010. **107**(13): p. 5881-5886.
4. Blazewicz, S.J., et al., *Evaluating rRNA as an indicator of microbial activity in environmental communities: limitations and uses.* The ISME Journal, 2013. **7**(11): p. 2061-2068.
5. Kozich, J.J., et al., *Development of a dual-index sequencing strategy and curation pipeline for analyzing amplicon sequence data on the MiSeq Illumina sequencing platform.* Applied and environmental microbiology, 2013: p. AEM. 01043-13.
6. Caporaso, J.G., et al., *QIIME allows analysis of high-throughput community sequencing data.* Nature Methods, 2010. **7**: p. 335.
7. Callahan, B., et al., *Bioconductor Workflow for Microbiome Data Analysis: from raw reads to community analyses [version 2; referees: 3 approved].* F1000Research, 2016. **5**(1492).
8. Callahan, B.J., et al., *DADA2: high-resolution sample inference from Illumina amplicon data.* Nat Methods, 2016. **13**.
9. Quast, C., et al., *The SILVA ribosomal RNA gene database project: improved data processing and web-based tools.* Nucleic Acids Research, 2013. **41**(D1): p. D590-D596.
10. Wang, Q., et al., *Naïve Bayesian Classifier for Rapid Assignment of rRNA Sequences into the New Bacterial Taxonomy.* Applied and Environmental Microbiology, 2007. **73**(16): p. 5261-5267.
11. Davis, N.M., et al., *Simple statistical identification and removal of contaminant sequences in marker-gene and metagenomics data.* Microbiome, 2018. **6**(1): p. 226.
12. McMurdie, P.J. and S. Holmes, *phyloseq: An R Package for Reproducible Interactive Analysis and Graphics of Microbiome Census Data.* PLOS ONE, 2013. **8**(4): p. e61217.
13. Swenson, V.A., et al., *Assessment and verification of commercially available pressure cookers for laboratory sterilization.* PLOS ONE, 2018. **13**(12): p. e0208769.
14. Lovley, D.R. and E.J.P. Phillips, *Rapid Assay for Microbially Reducible Ferric Iron in Aquatic Sediments.* Applied and Environmental Microbiology, 1987. **53**(7): p. 1536-1540.
15. Stookey, L.L., *Ferrozine---a new spectrophotometric reagent for iron.* Analytical Chemistry, 1970. **42**(7): p. 779-781.
16. Golterman, H.L. and R.S. Clymo, *Methods for chemical analysis of fresh waters*. 1969, Oxford; Edinburgh: published for the International Biological Programme by Blackwell Scientific.
17. APHA, *Standard methods for the examination of water & wastewater*. 2006, Washington, DC: American Public Health Association.
18. Ellis, P.S., et al., *Field measurement of nitrate in marine and estuarine waters with a flow analysis system utilizing on-line zinc reduction.* Talanta, 2011. **84**(1): p. 98-103.
19. APHA, *Standard methods for the examination of water and wastewater*. 1998, Washington, D.C.: APHA-AWWA-WEF.
20. Ioffe, B.V. and A.G. Vitenberg, *Head-Space Analysis and Related Methods in Gas Chromatography*. 1984, New York, USA: John Wiley and Sons.
21. R Core Team, *R: A language and environment for statistical computing*. 2018, R Foundation for Statistical Computing: Vienna, Austria.
22. RStudio Team, *RStudio: Integrated Development for R.* 2016, RStudio, Inc.: Boston, MA.
23. Oksanen, J., et al., *Package 'vegan'.* Community ecology package, version, 2013. **2**(9).

24. Knights, D., et al., *Bayesian community-wide culture-independent microbial source tracking.* Nature Methods, 2011. **8**: p. 761.
25. Pinheiro, J.C., et al., *nlme: Linear and Nonlinear Mixed Effects Models*. 2018.
26. Searle, S.R., F.M. Speed, and G.A. Milliken, *Population Marginal Means in the Linear Model: An Alternative to Least Squares Means.* The American Statistician, 1980. **34**(4): p. 216-221.
27. Lenth, R., et al., *emmeans: Estimated Marginal Means, aka Least-Squares Means*. 2019.
